# The *piggyBac* derived transposase 5 (PGBD5) can interact with human *piggyBac*-like elements

**DOI:** 10.1101/2025.07.31.667870

**Authors:** Linda Beauclair, Laura Helou, Florian Guilllou, Hugues Dardente, Thierry Lecomte, Alex Kentsis, Yves Bigot

**Author notes:** Joint Authors (Linda Beauclair, Laura Helou). Present Address: Linda Beauclair, Institut Jean-Pierre Bourgin, INRAE, AgroParisTech, Université Paris-Saclay,78000, Versailles, France; Laura Helou, Department of Human Genetics, CHU of Liege, Avenue de l’Hôpital 1, 4000, Liege, Belgium.

## Abstract

PGBD5 is encoded by a gene domesticated at the chordate origin from a DNA transposon of the *piggyBac* family. During its evolution, PGBD5’s sequence has been under strong purifying selection among vertebrate genomes. This suggests PGBD5 functions in the development and physiology of chordates, as recently demonstrated in mouse and human brain development, where it was implicated in double strand DNA breaks on neurons. However, biochemical PGBD5 activities remain undefined due to lack of appropriate in vitro model systems. Furthermore, coevolution of PGBD5 with hosts has likely reshaped some of its functions, resulting in differences between vertebrate PGBD5 and the insect *piggybac* transposase (PB). Recent studies have found that PGBD5 can interact with two different “species” of *piggyBac*-like transposon element (*pble*). Here, we show that human PGBD5 can interact with four “species” human *pbles* and to promote their chromosomal integration in cells, a property it shares with insect PB. Human PGBD5 can also bind to distinct chromosomal copies of human *pble* in cell type-specific manner, and to genomic loci containing inverted repeats in human cells akin to those found in subterminal insect *pble* ends. These findings expand the scope of potential biological activities of PGBD5 and other domesticated DNA transposases.

## Introduction

The gene *pgbd5*, stemming from a domesticated DNA transposon in the *piggyBac* family, stands out as one of the most evolutionarily conserved genes among vertebrates (1). Its origins date back approximately 500 million years, emerging from the domestication of a gene that encoded a transposase within a *piggyBac*-like element (*pble*), closely related to *Tcr-pble* (2). *Pble*s belong to the *piggyBac* family of eukaryotic transposable elements (TEs), utilizing a cut-and-paste transposition mechanism involving DNA as an intermediate (3,4). They are widely distributed throughout the genomes of various eukaryotic lineages (3). The first identified *pble*, *Ifp2* (5), an autonomous element spanning 2475 base pairs (bp), is commonly referred to as *piggyBac*. Originating from the genome of the cabbage looper moth (*Trichoplusia ni*), *Ifp2* serves as the reference element for *piggyBac* family research. *Ifl2*, along with other family members, inserts into chromosomes at the tetranucleotide TTAA sequence, which is duplicated upon insertion (4,6). These elements are flanked by terminal inverted repeats (TIRs) approximately 13 base pairs in length. Depending on the *pble* “species,” they may also exhibit subterminal inverted repeats (STIRs) with identical sequences or inner inverted repeats (IIRs) at both subterminal regions with differing sequences (2). In *Ifp2*, STIRs are 19 bp long and are internally located, positioned at 3 and 31 nucleotides from the 5’ and 3’ TIR inner ends, respectively (7,8). Within the inner region of the element, an open reading frame (ORF) about 1.8 kilobases in length encodes a transposase comprising approximately 600 amino acids (594 amino acids in the *Ifp2* transposase, also known as PB).

Multiple “species” of *pbles* underwent domestication at various points during the evolutionary history of different chordate lineages, giving rise to host *pgbd* genes encoding *piggyBac*-derived transposases (PGBD) (2,7). In addition to the *pgbd5* gene, the *pgbd* genes 1, 2, 3, and 4 are present in all simian species, resulting from four apparent domestication events at different times of Theria evolution (7). These genes encode the proteins PGBD1, PGBD2, PGBD3, and PGBD4.

Until recently, RAG1/2 was considered the sole domesticated transposase in vertebrate genomes, retaining most of its transposase activities and functioning as a site-specific recombinase (9). However, recent research has revealed that PGBD5 also retains transposase-like activities in cellular assays. Our investigations have demonstrated that PGBD5 enhances the genomic integration rate of a resistance gene, NeoR, carried by a DNA plasmid when it is transfected into the genomes of HEK293, HeLa, and G401 human cells. This effect is preferentially observed when NeoR is flanked at both ends by the 5’ and 3’ terminal regions of *Ifp2* (10,11,12) or *Tcr-pble* (2). Intriguingly, the nucleic acid sequences of *Ifp2* and *Tcr-pble* do not exhibit similarity, despite sharing the motif TTAACCC at both ends, a motif not uncommon in human genomes (218,809 exact and non-overlapping motifs in the hg38 genome reference). Hence, we have proposed that the specificity of interactions between PGBD5 and these two *pbles* is not related to their primary sequences but rather to the secondary structure of their ends or chromatin factors (2).

Results obtained from integration assays made with PGBD5, however, were challenging to interpret for two primary reasons. Firstly, PGBD5 displays high cytotoxicity in HeLa cells (2,10,14), leading to apoptosis and/or mitotic cellular arrests in a significant fraction of transfected cells in which it is expressed. Consequently, the results of integration assays must be adjusted to account for the reduced clonogenic capacity of cells expressing PGBD5. Secondly, in comparison to a “wild-type” transposase such as PB, PGBD5 appears to have weak cellular integration activity and less precise cleavage activities at the ends of *pbles* and its insertion sites. These characteristics result in the majority of integration events lacking sequence signatures consistent with canonical *pble* transposition, such as precise TTAA target site duplications (TSDs), intact TIRs and subterminal regions at their outer ends, and no traces of the plasmid backbone flanking the *pble* ends from the donor plasmid carrying the NeoR cassette. Since these non-canonical events occur in greater numbers in the presence of PGBD5 and stem from mechanisms of genomic integration that do not involve canonical transposition or homologous recombination, we will henceforth refer to them in this article as “PGBD5-dependent integration”. These challenges, along with others, including current lack of biochemical assays of purified PGBD5, have contributed to a controversy surrounding the consistency of these results, particularly in two papers that were themselves in part divergent (15,16). For example, PGBD5 can increase the integration rate of a NeoR cassette when flanked by the terminal sequences of two human *pbles*, MER75 and MER85 (16), as well as those of *Ifp2* (10,11,12,17).

Our current study aims to investigate whether human and mouse PGBD5 can interact with one or more of the four *pbles* found in human genomes, as well as with specific chromosomal motifs bearing similarities to *pble* subterminal regions. To assess PGBD5’s ability to affect the integration rate of a NeoR cassette flanked by human *pble* ends, we conducted integration assays in HeLa cells using four different PGBD5 isoforms from mouse (Mm) and human (Hs) origins, Mm523/Hs524, Mm409, Hs455, and Hs554 (17), along with four human *pble* sources, MER75, MER75B, MER85, and Looper, and the invertebrate *pble* representatives, *Tcr-pble* and-or *Ifp2* for comparison. Analyses were conducted to verify the presence of integration events resulting from canonical transposition or transposase-dependent integration events. To determine the cellular location of PGBD5 in transfected cells and select cell lines, we generated murine anti-PGBD5 sera using DNA vaccination technology ICANtibodiesTM (In Cell Art, Nantes, France) and employed a control procedure as described (18,19). These sera enabled the detection of PGBD5 through immunofluorescence in cell lines and defined its binding profile via chromatin immunoprecipitation sequencing (ChIP-seq) in specific cell lines. Our findings reveal that PGBD5 can bind to various types of repeats in mammalian cells that share motifs akin to *pble* STIR and IIR. Collectively, these results demonstrate that PGBD5 can interact with all four human *pbles*, as well as with *Tcr-pble* and *Ifp2*, two invertebrate *pbles*. These interactions, however, exhibit variability among different *pbles* and are significantly influenced by the nuclear environment, chromosomal DNA configuration, cell type, and the regulation of PGBD5 cellular biochemical activities.

## Materials and Methods

### Integration assay

#### Expression plasmids for proteins

Plasmid constructs were obtained as described (2,10). The plasmids pCS2-Mm523, pCS2-Mm409, pCS2-Hs554 and pCS2-Hs455 encode 5Xmyc-tagged PGBD5 isoforms that are 523, 409, 554 and 455 amino acid residues in length from mice (Mm) or human (Hs) origins, respectively. Our ORF source of Hs455 was the plasmid pRecLV103-GFP-PGBD5 (Addgene plasmid 65409; RefSeq accession numbers NP_078830 and NM_024554). The ORF encoding Hs554 (corresponding to RefSeq and Uniprot accession numbers NP_001245240 and EAW69912, respectively) was synthesized by ATG:biosynthetics (Merzhausen, Germany). Mouse Mm523 and Mm409 orthologues correspond to Uniprot/RefSeq accession numbers D3YZI9.21/NP_741958.1 and D3YZI9.12/XP_006530867.1, respectively. pCS2-GFP was described in (20) and pCS2-PB, pCS2-PB.1-558, and pCS2-PB.NLS-1-558 in (2). Here, PB corresponded to the mPB version.

#### Plasmid donors of a NeoR cassette

pBSK-NeoR was used as a source of NeoR cassette (sv40 promoter - ORF encoding resistance protein to G418/kanamycin-sv40 polyadenylation signal) void of any transposon sequences. The plasmids pBS-*Ifp2*-NeoR and pGH-*Tcr-pble*-TIR5’-NeoR-TIR3’ (pGH-*Tcr-pble*-NeoR) were described in (10) and used as transposon sources in excision and toxicity assays. The plasmids pGH-MER75, pGH-MER75B, pGH-MER85 and pGH-Looper were synthesized by ATG:biosynthetics (Merzhausen, Germany). Their sequence is supplied in supplementary data 1. A NeoR cassette was cloned between *Eco*RI and *Bam*HI sites to make pGH-MER75-NeoR, pGH-MER75B-NeoR, pGH-MER85-NeoR and pGH-Looper-NeoR.

#### Integration assay in HeLa cells

Essays were monitored as described (20). Briefly, each sample of 10^5^ cells in a well of a 24-wells plaque assays was co-transfected with JetPEI (N/P = 5 ; Polyplustransfection, Illkirch-Graffenstaden, France) and 400 ng DNA plasmid and with equal amounts of donor of NeoR cassette included or not within a transposon and transposase sources (1:1 ratio). Two days post-transfection, each cell sample was transferred to a cell culture dish (100 mm diameter) and selected with a culture medium containing 800 μg/mL G418 sulfate (Eurobio Scientific, Les Ulis) for 15 days. After two washing with 1X saline phosphate buffer, cell clones were fixed and stained overnight with 70% EtOH, 0.5% methylene blue and colonies > 0.5 mm in diameter were counted. Experiments were performed at least twice in triplicate.

#### Integration assay in G401 cells

Assays were monitored as described (11). Two clonal G401 cell lines were used to define the activities of human endogenous PGBD5. The first line was lentivirally transduced to constitutively express specific shRNA suppressing the expression of Hs524 PGBD5 (12). The second line was modified as a control to constitutively express shRNA to target GFP which is not expressed, thereby preserving the endogenous expression of Hs524 PGBD5 (the RNA-seq analysis of SRR13444179 to SRR13444184 datasets from the PRJNA692456 Bioproject confirmed that only mRNA encoding Hs524 and-or Hs409 were present in G401 cells). Briefly, each sample of 10^5^ cells in a well of a 24-well plates of plaque assays was transfected with jetOptimus and 500 ng DNA plasmid pBSK-IFP2-TIR5’-NeoR-TIR3’ as recommended by the supplier (Polyplus-transfection, Illkirch-Graffenstaden). Two days post-transfection, each cell sample was transferred to a cell culture dish (100 mm diameter) and selected with a culture medium containing 2 mg/mL G418 sulfate (Eurobio Scientific, Les Ulis) for 15 days. After two washing with 1X saline phosphate buffer, cell clones were fixed and stained overnight with 70% EtOH-0.5% methylene blue and colonies > 0.5 mm in diameter were counted. Experiments were performed at least twice in triplicate.

Data analysis, Krustal-Wallis tests (two-tailed), and graphical representations were performed using Anastats tools (available at https://www.anastats.fr/) and the Prism 9 package (GraphPad Software Inc.). Due to the small sample size (4 or 6 experimental replicates per sample), Kruskal-Wallis non-parametric tests were applied with a significance threshold of p < 0.05.

### Recovery of integration sites

#### LAM-PCR and Illumina libraries

Integration assays were done to produce cell populations containing integrated copies of the donor transposon. Fifteen days post-transfection, about 500 Neo clones were harvested for MER75 and MER75B, 1000 for MER85 and 200 for Looper. Genomic DNA preparations were made using the DNeasy kit (Qiagen, Hilden, Germany). Linear amplification-mediated PCR (LAM-PCR) was performed to amplify the vector-genomic DNA junctions of *Ifp2* vectors as described (21). All PCR were done using the high fidelity Q5 DNA Polymerase (New England Biolabs, Ipswich, MA). For both approaches, 1 µg DNA was used for twice 50 rounds of linear amplification using a biotinylated primer anchored near one end of the NeoR cassette to enrich DNA species containing transposon-chromosomal DNA junctions (for sequences of (B)-NeoR 5’ and 3’ primers, see supplementary data 1b). One reaction was done per end. The single-stranded products were immobilized on streptadivin-coated magnetic beads (Dynabeads M-280 Streptavidin, Invitrogen, Carlsbad, CA). All subsequent steps were performed on the magnetic bead-bound DNA. Two washes with water followed each step. Second strand synthesis was performed with random hexamer primers (Roche, Basel, Switzerland) using Klenow DNA polymerase (New England Biolabs, Ipswich, MA). The double-stranded DNA was split in two batches and subjected to restriction digests with *DpnI* for the first one and *Pci*I, *Nco*I and *Bsp*HI for the second one using restriction enzymes. The DNA fragments with a CG-3’ or a CATG-3’ overhang ends were ligated to linkers displaying appropriate overhang ends and made from annealed oligonucleotides (supplementary data 1b).

To increase the specificity of the full process, an initial PCR was done using one biotinylated primer anchored within the 5’ or 3’ region of the transposon donor and one primer anchored within the linker (for sequences of (B)-TIR-UTR 5’ and 3’, and LC1 primers, see supplementary data 1b). PCR products were immobilized on streptavidin-coated magnetic beads and purified as described above. Next, the bead-bound DNA was subjected to a nested PCR using nested primers anchored within transposon ends and within linkers (supplementary data 1c). Final PCR products were purified, quantified and gathered in equimolar DNA amounts for each transposon vector (4 populations of LAM-PCR products) before being used to make Illumina libraries using NEBNext® Ultra™ II DNA Library Prep Kit for Illumina® and NEBNext Multiplex Oligos for Illumina (New England Biolabs, Ipswich, MA). Fragment size selection, library quality control and Illumina sequencing (MiSeq 250 nucleotides, TruSeq SBS Kit v3) were achieved at the Plateforme de Séquençage Haut Débit I2BC (Gif-sur-Yvette, France). DNA quantities were monitored at various steps in the procedure with the Qubit® dsDNA (Molecular Probes, Eugene, USA).

#### Computer analysis

Trimmomatic (22) was used to filter Miseq reads using default parameters, except for SLIDINGWINDOW:5:20 and MINLEN:100. The purpose of the following steps was to identify junctions between chromosomal DNA and inserted DNA fragments, taking into account the plasmid backbone regions located 100-bp upstream and downstream *Ifp2*-NeoR transposon. Filtered reads were first mapped to the sequence of plasmid backbone minus the 100-bp regions flanking both sides the *Ifp2*-NeoR transposon with bwa-mem using default parameters (23). Unmapped reads were then extracted using SAMtools view with parameters -b -f 4 (24), converted back to FASTQ format using bamt bamtofastq from the BEDTools suite using default parameters (25), and finally realigned to a custom reference containing hg38 chromosomes and the *Ifp2*-NeoR transposon flanked by the 100-bp plasmid backbone regions (supplementary data d), using BWA-MEM with modified parameters (-w 1 -r 1). The bam files resulting from each dataset alignment were analysed with Lumpy in order to identify split reads (26). The parameters were -e -mw 2 -tt 0.0 and back_distance:20,weight:1,id:lumpy_v1,min_mapping_threshold:20. Structural variants (SV) annotated as “BND” (breakend) and characterized by paired breakpoints, one located in the genome and the other within the transposon,-, were extracted using a house python script (https://github.com/Leelouh/lumpy2site). Results were filtered taking into account a difference below 3 between the transposon breakpoint calculated by Lumpy and the maximal spread of read alignments in the transposon donor sequence for each integration event. Each TSD nucleotide motif at the insertion site was obtained after extracting 10-bp sequences before and after the breakpoint in the chromosome sequences.

### RNA expression profiles in our HeLa cells

#### Culture of cell lines and stress treatment

HeLa (cervical carcinoma), HEK293T (kidney), HCT116 cells (colon), and H4 (brain epithelium) cell lineages were cultured in Dulbecco’s modified Eagle’s medium (DMEM) supplemented with 10% fetal bovine serum (FBS). Each stress treatment was monitored 24 hours after having seeded 2.5x10^6^ cells in each well of a 6-well plate of plaque assays that was maintained at 37°C and with 5% CO_2_. For the heat shock, cells were incubated for 3 hours at 42°C and with 5% CO_2_. For chemical treatment, the medium was adjusted at 500 nM in geldanamycin and 100 nM radicicol (ENZO Life Sciences, Villeurbanne, France) and cells were maintained for 1 hour at 37°C and with 5% CO_2_. At the end of each of these treatments, cells were trypsinized, washed twice with 1XPBS and total RNA were finally purified using the NucleoSpin Extract RNA kit (Macherey Nagel, Hœrdt, France).

#### Illumina library and sequencing

Libraries for Illumina sequencing were made using NEBNext® Ultra™ II RNA Library Prep Kit for Illumina® (New England Biolabs, Ipswich, MA, USA). DNA quantities were monitored at various steps in the procedure with the Qubit® dsDNA HS Assay Kit (Molecular Probes, Eugene, USA). Fragment size selection, library quality control and Illumina sequencing (HiSeq 51 and 76 nucleotides, TruSeq SBS Kit v3) were achieved at the Plateforme de Séquençage Haut Débit I2BC (Gif-sur-Yvette, France), following published quality recommendations (27). Data published in this paper are based on two biological replicates.

#### Computational analyses of RNA-Seq

Datasets were filtered using FastQC and Trimmomatics. Each set of reads was mapped to hg38 using RNA-STAR (28). Read alignments were visualized with IGV, quantified with featureCounts and genes with either increased or decreased expression were detected using DEseq2 and edgeR (29,30).

### Profile of PGBD5 binding in several human cell lineages

#### Culture of cell lines

HeLa (uterus), HEK293T (kidney), HCT116 cells (colon), and H4 (brain epithelium) cell lineages were cultured in Dulbecco’s modified Eagle’s medium (DMEM) supplemented with 10% FBS. T98G (glioblastoma) and 8-MG-BA (glioblastoma) cell lineages were respectively cultured in minimum essential medium Eagle with Earle’s salts (MEM), supplemented with 10% FBS and 1mM L-glutamine or RPMI 40%, MEM 40%, 20% FBS. The six cell lineages were maintained at 37 °C and with 5% CO_2_. The sources of the cell lineages were those of the EA GICC 7501 (CHRU de Tours, 37044 TOURS Cedex 09) which were acquired from ATCC (HeLa (ATCC® CCL-2); HEK293T (ATCC® CRL-3519); HCT116 (ATCC® CCL-247EMT); H4 (ATCC® HTB-148); T98G (ATCC® CRL-1690), 8MGBA (DSMZ ACC432).

#### Antibodies

As previously reported (18), the quality of antibodies is crucial to study the expression of domesticated transposases. Because there is no suitable anti-PGBD5 commercial antibody for ChIP, western blot and immunohistochemical analyses, the anti-PGBD5 referenced orb13159 (Biorbyt Ltd, Cambridge, UK) being no longer distributed since 2017, the alternative products being unsatisfactory (orb389313 and orb620969) like other anti-PGBD5 antibodies commercially distributed (including the M1012.1 monoclonal antibody from HUABIO), murine sera containing polyclonal antibodies (pA) were produced using DNA vaccination technology ICANtibodies^TM^ (In Cell Art, Nantes, France). These sera were obtained using pVAX mammal expression plasmid coding the Mm409 isoform of mouse Pgbd5. Such sera have been reported to be effective for Western blots, ELISA, cells spread on slide and ChIP (18,19). Five SWISS and five C57/B6 male mice were injected at day 0, 21, 42 and 56. Blood samples were harvested before the first injection and at days 21, 42 and 56. The complete blood was recovered at day 71. The presence of anti-PGBD5 pA directed against linear and 3D epitopes was verified (see result section). The protein concentration in the final serum was defined using the BCA Protein quantification Kit (Interchim, Montluçon, France).

#### ChIP-Seq experiments

Chromatin samples were prepared from non-synchronous and exponentially growing cells. Chromatin shearing was performed with a Bioruptor ultrasonicator (Diagenode, Ougrée, Belgium). Chromatin immunoprecipitation was performed with 10 µL of pre-immune or anti-PGBD5 sera, and purification of immunoprecipitated DNA was done using the iDeal ChIP-Seq kit following the supplier’s recommendations (Diagenode). Libraries for Illumina sequencing were made using NEBNext Ultra II DNA Library Prep Kit for Illumina (New England Biolabs, Ipswich, MA, USA). DNA quantities were monitored at various steps in the procedure with the Qubit® dsDNA HS Assay Kit (Molecular Probes, Eugene, USA). Fragment size selection, library quality control and Illumina sequencing (HiSeq 51 nucleotides, TruSeq SBS Kit v3) were achieved at the Plateforme de Séquençage Haut Débit I2BC (Gif-sur-Yvette, France), following published quality recommendations (27). Data published in this paper are based on at least three biological replicates.

For HeLa, HEK293T, HCT116, H4, T98G and 8-MG-BA cell lineages, sequence reads were mapped to the human genome assembly hg38 (December 2013; available at http://www.ncbi.nlm.nih.gov/assembly/GCF_000001405.26) with the Bowtie short read aligner (31). Peak calling was done with two tools from bam files and normalized over input (three biological replicates), the peak-calling prioritization pipeline PePr1.1.16 (32). For both tools, a q-value threshold of 5.10^-2^ was used. Other parameters were those per default for PePr1.1.16. peaks. For the G401 cell lines, data from NCBI Sequence Read Archive PRJEB41045 and PRJEB41052 projects were reanalysed using the above procedure. The peak number and their locations were similar to those previously described (11). Peaks of each cell lineage were filtered using the ENCODE blacklist (33). Intersections between ChIP-Seq peaks and the other annotation files were calculated using bedtools.

#### Computational analyses of ChIPseq peaks

Annotations of human *pble* fragments in hg38 were extracted from the RepeatMasker (RM) annotations as starting points. These annotations were then completed as described in (19). For MER75, there were 801 annotations in RM, 1211 in ours, for MER85, 929 versus 1067, and Looper, 559 versus 1362. This was mainly due to the fact that numerous human pbles were fragmented but part of these fragments was not annotated in the RM annotation. The annotation of *pble*-like inner inverted repeats was done using Palindrome (EMBOSS package). The output file was then filtered using pal2gff (https://github.com/Leelouh/pal2gff/blob/main/pal2gff.py), using as parameters a repeat size between 5 and 15 nucleotides, a spacer between pairs of inverted repeats (IRs) of 2 to 10 nucleotides, and a number of mismatches within repeats ranging from 0 to 1. These parameters were chosen taking into account those of the inner IRs found at ends of invertebrate *pbles* (2). The final file contained 28,502,545 annotations and is available at https://doi.org/10.5281/zenodo.8281129. Searches for conserved motifs were carried out with the MEME suite (MEME-ChIP and GLAM2) (34) and the RSAT pipeline (35,36).

Data analyses, t-tests (two-tailed) and graphic representation were done using bedtools and Prism 9 package (GraphPad Software Inc). Before t-tests, a F test was done to compare variances of samples and a Welch’s correction was used when they were significantly different (p>0.05). Permutation tests (10,000 per test) were computed using shuffleBed (with options - noOverlapping and -chromFirst) to produce samples containing non-overlapping features and intersectBed to count overlaps between two feature files. Probabilities were calculated from Z score at https://www.fourmilab.ch/rpkp/experiments/analysis/zCalc.html.

### Cellular localization of green fluorescent protein-fusion proteins

#### Plasmid expression for transposase-GFP fusions

The plasmids pCS2-GFP, and pCS2-Mm523-GFP were made as described (37). The plasmids pCS2-Mm523 and pCS2-Mm409 were described above.

#### Cell manipulation

HeLa cells were plated at a density of 5 x 10^4^ cells per well in 1 cm^2^ Lab-Tek^TM^ chamber slides (Fisher Scientific, Waltham, MA, USA) and grown in DMEM (Gibco/Life Technologies, Paisley, UK) supplemented with 10 % heat inactivated fetal bovine serum (FBS, Eurobio, France) at 37 °C in a humidified atmosphere containing 5% CO2 for 48 h. Cells were transfected with 500 ng plasmid DNA and jetPEI™ (Polyplus Transfection, Illkirch, France) at a N/P ratio of 5 (basically the ratio of positively-chargeable polymer amine (N = nitrogen) groups to negatively-charged nucleic acid phosphate (P) groups) in DMEM 10% FBS following the manufacturer’s instructions. Cells were then incubated with the complexes for 4 h. The transfection medium was then discarded and replaced by fresh DMEM supplemented with 10% FBS before being incubated for 48 hours at 37°C.

#### GFP Imaging

Two days post-transfection, cells on slides were washed three times with 1X-PBS at room temperature (RT), then fixed in 1X PBS/2% paraformaldehyde at RT for 15 min, and then permeabilized with PBS/1% (w/v) Triton-X100 for 10 min. The slides were washed three times for 5 min with 1x PBS. Nuclei were stained using Vectashield Vibrance “Antifade Mounting Medium (hardening) + DAPI” (Vector Laboratories, Burlingame CA, USA). All images of fluorescence were collected with an LSM 700 laser scanning microscope and the associated Zen software (Carl Zeiss, Oberkochen, Germany).

#### Imaging after antibody staining

Two days after transfection or plating, the cells were washed three times with 1X-PBS at room temperature (RT), cleaned with 1X-PBS, 5 µg/ml saponin for 3 min at RT, and then fixed with 3.7% paraformaldehyde for 30 min at RT. After fixation, the cells were washed three times for 5 min with 1X-PBS at RT, before being permeabilized in 1X-PBS, 0.5% Triton X100 for 5 min at RT, and then washed three times for 5 min at RT with 1X-PBS, 2% bovine serum albumin (BSA). Cells were incubated for 1 h at room temperature with a 1/100 dilution of a mouse antiserum (directed against PGBD5) in 1X-PBS, 2% BSA. The cells were then washed three times for 5 min with 1X-PBS, 0.2% Tween20 before being incubated for 1h, at RT with the secondary antibody (Donkey anti-Mouse CY3, Beckman Coulter). The slides were washed three times for 5 min with 1X-PBS, 0.2% Tween20. Nuclei were stained using Vectashield Vibrance “Antifade Mounting Medium (hardening) + DAPI” (Vector Laboratories, Burlingame CA, USA). All images of fluorescence were collected with an LSM 700 laser scanning microscope and the associated Zen software.

#### Confocal pictures

All images shown correspond to one focal plane (0.5 µm). Images to be used for figures were pseudocolored by LSM Image browser software (Carl Zeiss, Thornwood, NY). Photoshop (Adobe Systems, San Jose, CA) and ImageJ 1.46r were used on the resulting tiff files only to adjust for brightness and contrast, and to make assembly with Z-stack images.

### PAGE, immunoblotting and hybridization of antisera

#### Production of protein extracts

To mimic conditions in transposition assays used in the literature, HeLa cells were transfected as described in the above section “Integration assay in HeLa cells” using 400 ng (10,20) DNA plasmid and with equal amounts of pBS-*Ifp2*-NeoR (2,10) and pCS2-Tranposase (1:1 ratio; pCS2-PB, -PB.1-558, -PB.NLS-1-558, -Mm523, -Mm409, - Hs554, -Hs455 or pBS-SK+ as a negative control) and JetPEI (N/P = 5 ; Polyplustransfection, Illkirch-Graffenstaden, France).

Because 48 hours post transfection, floating dead cells in the middle of the well in assays done with all proteins were replaced by adherent non-transfected cells on the bottom of the well, analyses were performed at 24 hours post transfection, i.e. after about 15 hours of measurable protein expression (20). After two washings in PBS1X, proteins in adherent cells were extracted by vigorous pipetting with 300 µL 1X Laemmli buffer, 0.1% 2-Mercaptoethanol at room temperature (RT), sonicated three times 15 seconds at 40 W, 46 kHz using an Ultrasonic Jewelry Cleaner Leo (Leo, New Taipei City, Taiwan). Protein samples were then heated at 95°C for 10 min, cooled 5 min at RT and centrifuge 1 min at 15000 g at RT.

#### Rate analyses of double-strand DNA breaks and cell number

The induction of DNA damage repair in a cell population transfected with a Tranposase plasmid source was measured using phosphorylated γ-H2AX (phosphorylation of the Ser-139 residue of the histone variant H2AX) in each cellular protein extract (38). Samples (20 µL) were resolved by electrophoresis on a 4-15% mini-Protean TGX gel done in Tris/Glycine/SDS buffer (Bio-Rad, Hercules, USA). Gels were then transferred onto 0.2 µm nitrocellulose membranes using a Trans-Blot Turbo Transfer System gel (Bio-Rad, Hercules, CA, USA). Membranes were probed in Odyssey blocking buffer, 0.1 % Tween 20 at 4°C with a mouse anti-phospho-H2A.X (Ref. 9718, Cell Signalling Technologies, Danvers, MA, USA), a rabbit polyclonal anti-PRPF19 (Ref. PA5-24797, Invitrogen, Waltham, MA, USA) and a mouse anti-tubulin (Ref. T9026, Sigma, Saint-Louis, MI, USA), all in a dilution of 1/1000. After three washings in PBS1X, 0.1 % Tween 20 at RT, membranes were probed for 1 hour at RT in Odyssey blocking buffer, 0.1 % Tween 20 with secondary antibodies, an anti-mouse conjugated to IR-800 dye and an anti-rabbit conjugated to IR-670 dye, both at a 1/10000 dilution. After three washings in PBS1X, 0.1 % Tween 20 at RT, γ-H2AX, α-tubulin and PRPF19 were detected by near-infrared fluorometry, using an Odyssey scanner (LICOR Biosciences, Lincoln, NE, USA). Fold changes in amounts of H2A.X phosphorylation and PRPF19 were calculated by normalizing the band intensities with tubulin band intensities using Odyssey scanner facilities. Five replicates were done for each sample from the transfection to the Western blot analysis.

#### Detection of anti-PGBD5 antibodies in ICA mouse sera

The detection of 5Xmyc-Mm523 PGBD5 isoform in transfected HeLa cells was done using a similar procedure in which the primary antibody used for detection was a mouse anti-c-myc monoclonal antibody (Ref. 11 667 149 007, Roche, Bâle, Switzerland) at 1/1000 dilution.

### Manuscript preparation

The corresponding author was assisted by DeepL and Antidote 11 (Druide informatique inc, Montréal, Canada) to improve readability, which was thereafter optimized using ChatGTP. The final manuscript was edited manually by all authors.

## Results

### PGBD5 exhibits active genomic integration of a NeoR cassette flanked by human *pble* ends in human cells

To explore PGBD5’s capability to integrate a NeoR cassette flanked by human *pble* ends into mammalian cell chromosomes, we employed two distinct integration assays. The first assay was conducted in HeLa cells, which were transiently transfected with two plasmids: one carrying the transposon (pGH-MER75-NeoR, pGH-MER75B-NeoR, pGH-MER85-NeoR, pGH-Looper-NeoR, and pBSK-*Tcr-pble*-NeoR, see supplementary data 1), and the other providing the PGBD5 isoforms, Mm523, Mm409, Hs455, and Hs554. GFP was used as a negative control, and the expression of all four PGBD5 isoforms and GFP was driven by a CMV promoter. Under the experimental conditions used in this assay, we observed a reduction in the number of cells 24-hours post transfection in the presence of all PGBD5 isoforms, except Hs455, Hs554 and the PB.1-558 mutant (Figure S1A). Furthermore, an increase in phosphorylated γH2AX, a biochemical surrogate of DNA damage repair, was observed in cells transfected with plasmids expressing the PGBD5 isoforms, except Hs455, Hs554 and the two PB.1-558 mutants (Figure S1B). This indicated that many transfected cells underwent cell death or growth arrest, confirming previous observations of PGBD5 cytotoxicity (2,10,14,17). Despite these issues, we were still able to obtain NeoR clones, albeit at a lower rate compared to the control with GFP as the protein source. The raw results were depicted in graphs A, C, E, G, and I of Figure 1. To account for PGBD5-induced reductions in clonogenic capacity (2,10,17), estimated to be approximately 12-, 11-, 3-, and 2-fold for Mm523, Mm409, Hs455, and Hs554, respectively (2,17), we adjusted the results in graphs B, D, F, H, and J of Figure 1.

**Figure 1.**
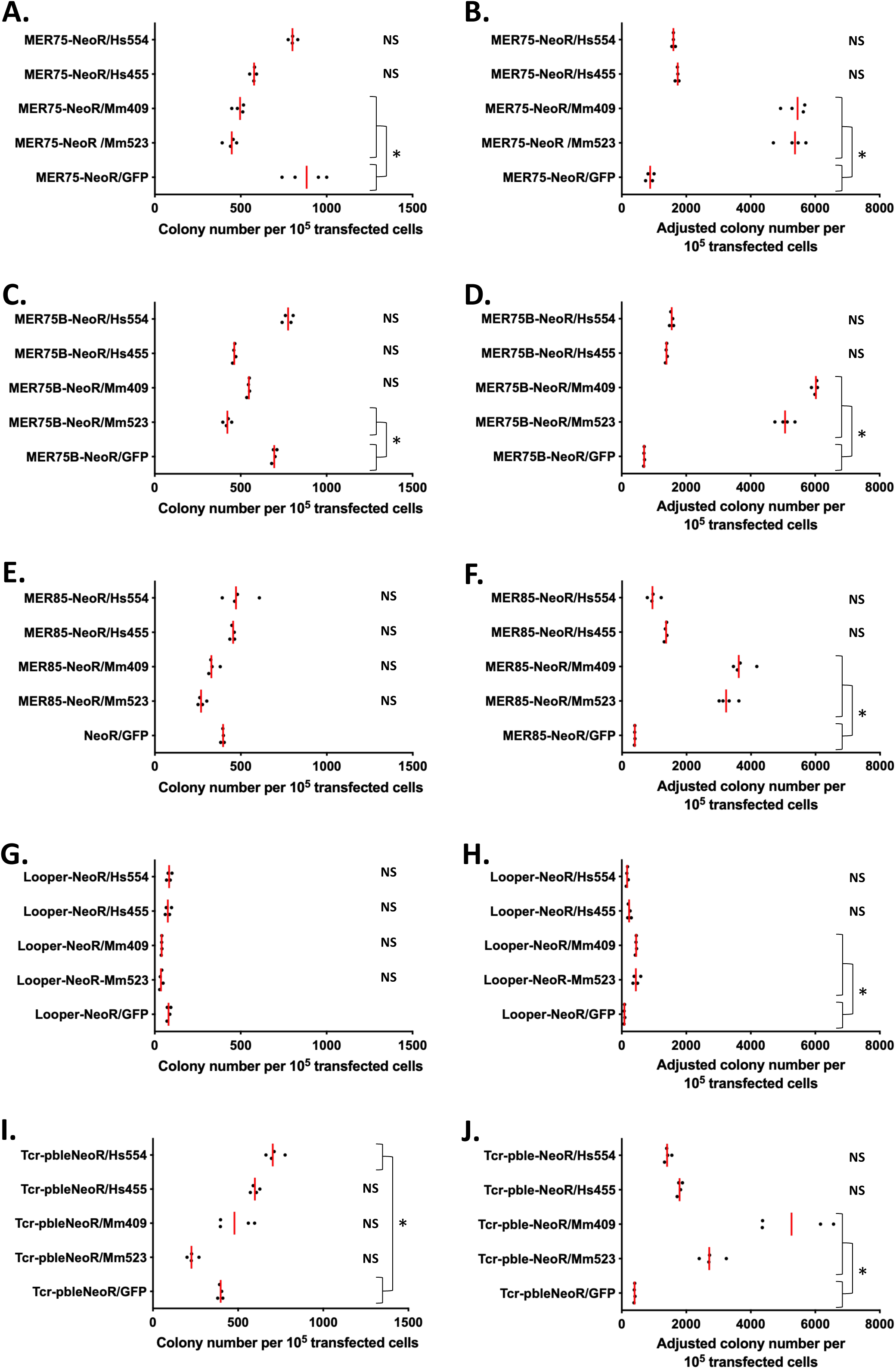
Clonogenic capacity of NeoR cassette flanked by *pble* ends to integrate into the genome of HeLa cells. Numbers of NeoR clones resulting from the chromosomal integration of a NeoR cassette flanked by MER75 (**A**), MER75B (**C**), MER85 (**E**), Looper (**G**) or *Tcr-pble* (**I**) ends in presence of GFP, Mm523, Mm409, Hs455 or Hs554. Numbers of NeoR clones resulting from the chromosomal integration of a NeoR cassette flanked by MER75 (**B**), MER75B (**D**), MER85 (**F**), Looper (**H**) or *Tcr-pble* (**J**) ends in presence of GFP, Mm523, Mm409, Hs455 or Hs554 once the values were adjusted by the impairment of NeoR clonogenic capacity measured in (17). Median values are marked by red lines and were calculated from 4 replicates. * and brackets, and NS indicated a significant and a non-significant Krustal-Wallis test (p>0.05) between results obtained with Mm523, Mm409, Hs455 or Hs554, and GFP, respectively.

Adjusted results demonstrated that the presence of two PGBD5 isoforms (Mm523 and Mm409) significantly increased the number of NeoR clones, indicating a higher integration rate of DNA fragments containing NeoR from plasmids pGH-MER75-NeoR, pGH-MER75B-NeoR, pGH-MER85-NeoR, pGH-Looper-NeoR, and pBSK-*Tcr-pble*-NeoR. Interestingly, it was noted that the two isoforms with the greatest changes in clonogenic capacity changes were also the most effective in enhancing the chromosomal integration of the NeoR cassette. Among the transposon DNA substrates, Looper seemed to be a DNA substrate less favorable for PGBD5 isoforms. Regarding the activity rates of these isoforms, it is essential to consider that the protein concentrations of each isoform in HeLa cells were different, with normalized values reported as 1 for Mm524, 0.39 for Mm409, 0.57 for Hs455, and 0.17 for Hs554 (17).

The second assay involved transfecting a NeoR plasmid donor, flanked by *pble* ends, into two human rhabdoid tumor G401 clonal lines. One of both G401 clonal line constitutively expressed endogenous Hs524 (12), referred to as G401-shGFP in Figure 2, which expresses a control short-hairpin RNA (shRNA) targeting non-expressed GFP. The second clonal line has its endogenous Hs524 expression reduced due to the expression of a constitutive shRNA specifically targeting Hs524 mRNA. It was referred to as G401-shPGBD5 in Figure 2. We confirmed reduction of endogenous PGBD5 using quantitative reverse transcriptase PCR (qRT-PCR; 11,12). NeoR clones were obtained for all assayed *pbles*, except for Looper in which no statistically significant differences between both cell lines were detected (p=0.13), and for the negative control (p=0.33). These results further confirmed PGBD5’s ability to interact with various *pbles*, with *Tcr-pble* being a more suitable substrate than MER75, MER75B, and MER85 for integrating a NeoR cassette, while Looper was not conducive to the integration. Integration rates differed significantly between *Tcr-pble* and MER75, MER75B, and MER85. However, it should be noted that the experimental setups in these two sets of experiments were considerably different, potentially contributing to the variations in PGBD5 activity, especially regarding its expression differences between transiently transfected HeLa cells and endogenously-expressing G401 cells.

**Figure 2.**
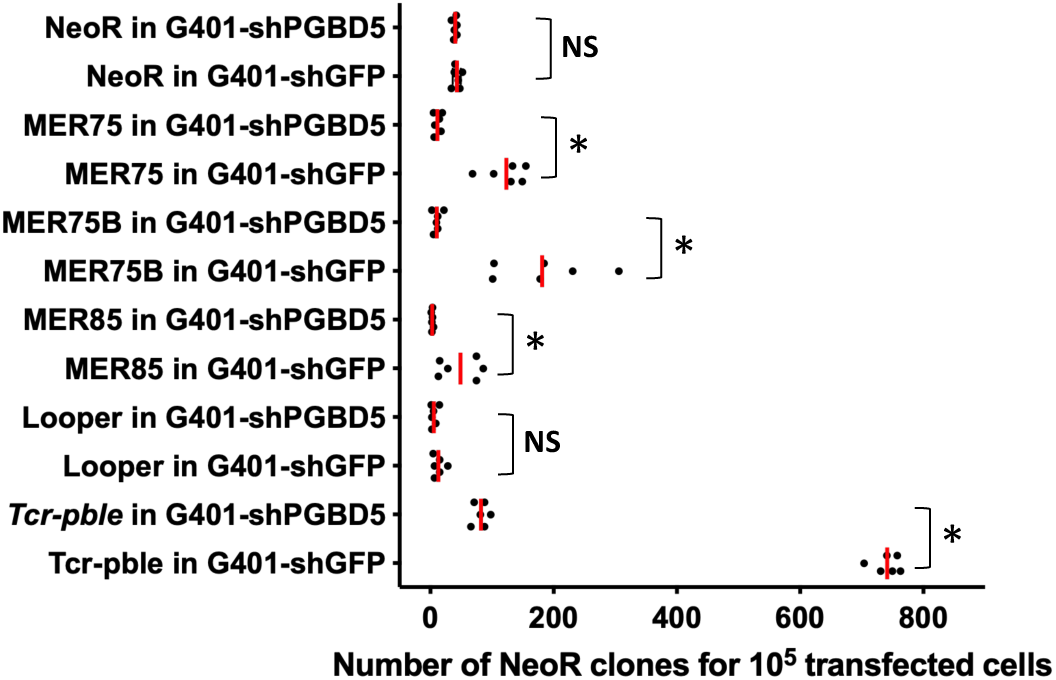
Clonogenic capacity of NeoR cassette flanked by *pble* ends to integration into genome of G401 cells expressing or not the PGBD5 isoform Hs524. Number of NeoR clones resulting from the chromosomal integration of a NeoR cassette alone (NeoR) or flanked by MER75, MER75B, MER85, Looper, Ifp2 or *Tcr-pble* ends in presence of PGBD5 (G401-shGFP cells) or in absence (G401-shPGBD5 cells). Median values are marked by red lines and were calculated from 2 x 3 replicates. * and brackets, and NS indicated a significant and a non-significant Krustal-Wallis test (p>0.05) between G401-shGFP and G401-shPGBD5 cells for each transposon source, respectively. For MER85, p=0.0198. The PGBD5 expression profiles of both clonal lines was validated in (11) and its low rate in G401-shGFP in (14).

### PB transposase also enhances the integration of human PBLE vectors

Given that PB transposase has been shown to bind to murine ES cell chromosomal DNA, possibly depending on the distribution of the tetranucleotide TTAA (39), and because it can mobilize distantly related *pbles* like *NlPLE25* (40), we explored whether PB could increase the integration of a NeoR cassette flanked by other *pble* ends. Integration assays in HeLa cells were conducted using various pbles as transposon sources (pGH-MER75-NeoR, pGH-MER75B-NeoR, pGH-MER85-NeoR, pGH-Looper-NeoR, and pBSK-*Tcr-pble*-NeoR), with pBSK-NeoR serving as a negative control, and pCS2-PB and pCS2-GFP as a source or not of PB, respectively. Results (Figure 3) indicated that the integration rates of *pbles* into chromosomes were significantly higher (approximately 2-fold) than in the GFP control for all assayed *pbles*, except for Looper, which showed no difference from the control (p=0.054). However, it’s worth noting that the control with pBS-NeoR yielded fewer NeoR clones in the presence of PB than in its absence, albeit weakly (1.15 to 1.30-fold). This difference was statistically significant, as observed in previous results (10,17). To conclude, these findings corroborated that PB has the ability to bind to DNA sequences beyond *Ifp2* ends and shares with PGBD5 the capacity to enhance the integration of a NeoR cassette flanked by *pble* ends other than those of *Ifp2*.

**Figure 3.**
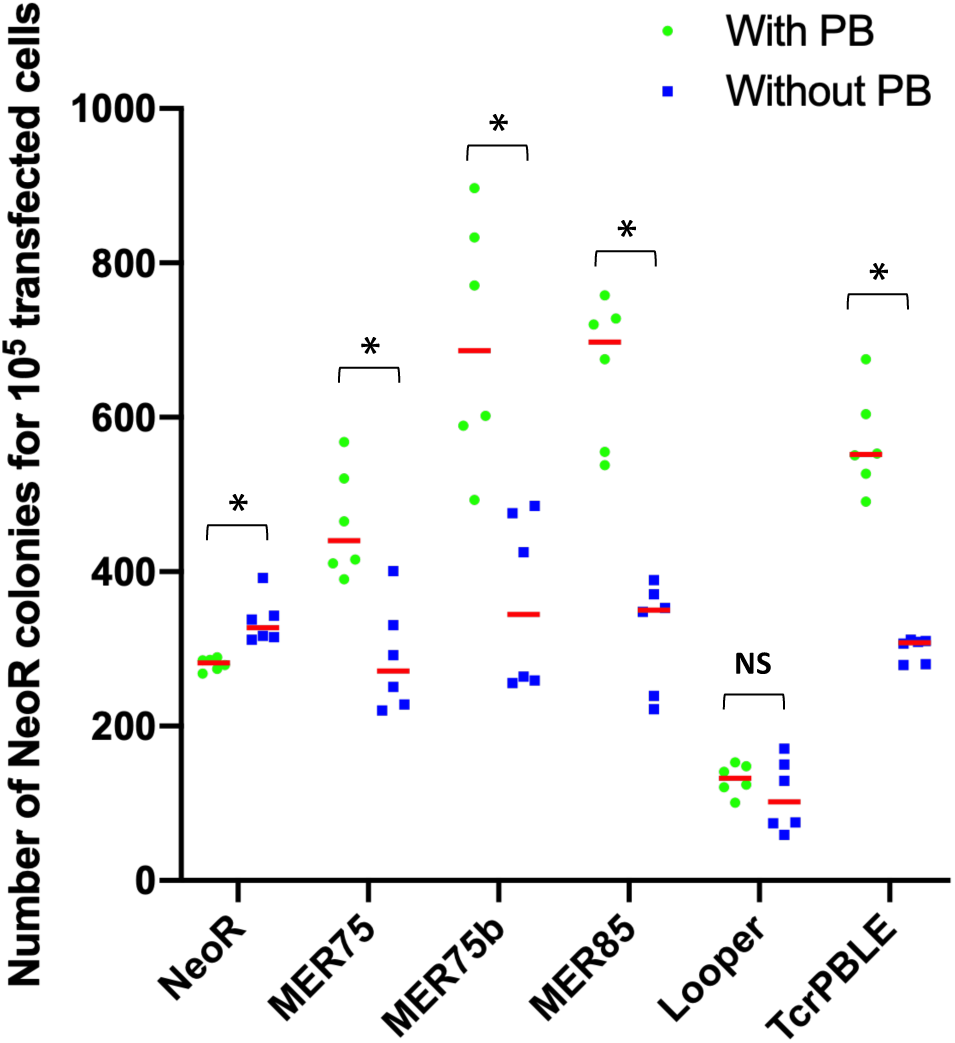
Clonogenic capacity of NeoR cassette flanked by *pble* ends to integrate into the genome of HeLa cells expressing or not the PB transposase. Number of NeoR clones resulting from the chromosomal integration of a NeoR cassette flanked by MER75, MER75B, MER85, Looper or *Tcr-pble* ends or with no flanking *pble* sequences (pBS) in presence or in its absence (the PB transposase). Median values are marked by red lines and were calculated from 2 x 3 replicates. * and brackets, and NS indicated a significant and a non-significant Krustal-Wallis test (p>0.05) between cells expressing or not PB for each transposon source. For pBS, p=0.001364.

### Features of human *pble* ends integrated into chromosomes

The process of PGBD5 integration unfolds as follows: a NeoR cassette, flanked by human *pble* ends, becomes integrated. To delve into the intricate features of these integration sites, we generated fragment populations corresponding to MER75-, MER75B-, MER85-, and Looper-chromosome junctions. These populations were derived from LAM-PCR amplifications utilizing genomic DNA (gDNA) extracted from NeoR clone populations. The sequencing of these populations was carried out employing Illumina Miseq technology. For gDNA sample preparation, approximately 500 clones were used for integration assays involving MER75 and MER75B, around 1000 clones for MER85, and roughly 200 clones for Looper. Our previous findings, drawn from integration assays conducted in HeLa cells (2,10), indicated that when PB served as the transposase source, the rate of *Ifp2* integration into chromosomes via proper transposition, marked by a perfect “TTAA” TSD and TIR sequence, exceeded 96%. This rate dropped to 19% when employing PB-NLS.1-558, a PB variant devoid of the C-terminal like PGBD5, and further to 12.9% with Mm523 PGBD5.

We detected 383, 306, 1313, and 121 unique junctions for MER75, MER75B, MER85, and Looper, respectively (Figure 4, supplementary data 2). Echoing our previous findings with *Ifp2* and *Tcr-pble*, we observed that the majority of these junctions did not result from proper transposition events (Table 1). Specifically, MER75 and Looper exhibited rates 3.6- and 1.9-fold higher than expected for proper TSD and TIR events, suggesting that at least some of these junctions might be interpreted as resulting from proper transposition, at least at one of their ends. Differing from our observations with *Ifp2* and *Tcr-pble* (2), most junctions were situated within the plasmid backbone sequences flanking each *pble*, rather than within the transposon ends.

**Figure 4.**
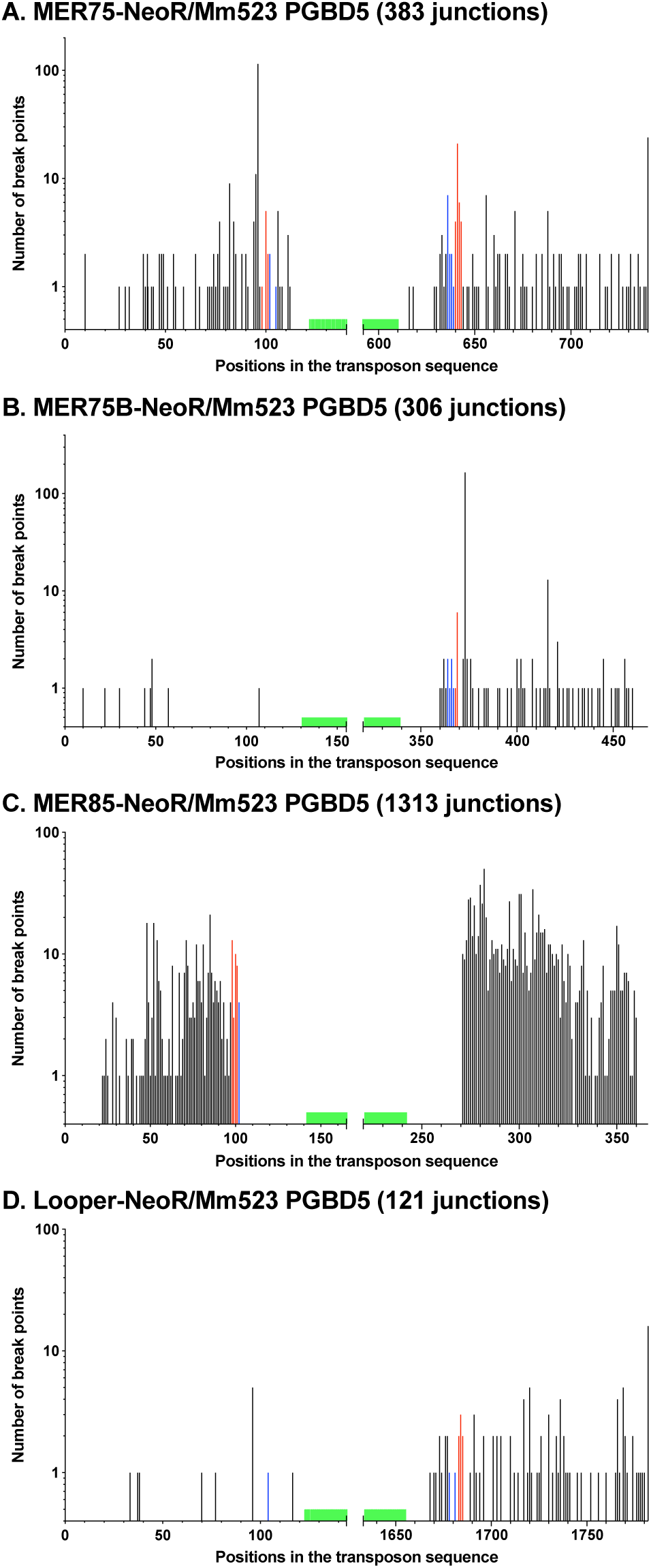
Number and location of transposon breakpoints in human *pble* sequences after integration into chromosomes. Histogram distributions of *pble*-NeoR extremities integrated by Mm523 (supplementary data 2). Red bars indicated insertion events with perfectly conserved TSD and TIR while blue bars located those in which TIR were perfectly conserved but the TSD did not correspond to a canonical TTAA at the outermost extremities of *pble*s. Black bars represented breakpoints within the transposon sequence and within plasmid backbone sequences juxtaposed to the transposon. Each bar corresponded to the number of junctions found in a single nucleotide position. Green boxes located the position of primers anchored within the transposon sequence and used at the last step of LAM-PCR. These graphics described the relative importance of wounds at transposon ends under our experimental conditions. However, they could not allow calculating wound rates at each of both ends due to the fact that the final LAM-PCR products in each dataset came from the gathering of two independent LAM-PCR reactions done at the 5’ and 3’ ends.

**Table 1.** Presence of canonic TTAA TSD and-or TIR among transposon/chromosome junctions resulting from LAM-PCR products and originating from events mediated by 2 PB variants or the PGBD5 isoform, Mm523.

| Transposase source | Transposon source | % Proper TSD and TIR* | % Proper TSD improper TIR** | % Improper TSD/proper TIR** | % Improper TSD and TIR** |
| --- | --- | --- | --- | --- | --- |
| PB (7) | *lfp2*-NeoR | 96.7 (2.7%; 7370) | 1.05 (78) | 1.05 (80) | 1.2 (95) |
| PB.NLS-1-558 (7) | *lfp2*-NeoR | 19.0 (2.7%; 98) | 2.5 (13) | 3.1 (16) | 75.4 (3.8%) |
| Mm523 (8) | *lfp2*-NeoR | 12,9 (2.7%; 147) | 5.4 (79) | 6,4 (93) | 75.3 (1142) |
| Mm523 (8) | *Tcr-pble*-NeoR | 3.4 (2.7%; 60) | 5.7 (101) | 3.7 (66) | 87.2 (1539) |
| Mm523 (this study) | MER75-NeoR | 11.2 (3.1%; 43) | 1.6 (6) | 3.9 (15) | 83.3 (319) |
| Mm523 (this study) | MER75B-NeoR | 2.3 (3.1%; 7) | 3.9 (12) | 2.0 (6) | 91.8 (281) |
| Mm523 (this study) | MER85-NeoR | 2.5 (3.1%; 33) | 2.7 (36) | 0.3 (4) | 94.4 (1240) |
| Mm523 (this study) | Looper-NeoR | 5.8 (3.1%; 7) | 2.5 (2) | 2.5 (3) | 90.1 (109) |

\*, between parentheses are indicated i) the average percentage if the junctions were distributed at random within the window spanning from the use primer (in green in Figure 4) to about 100 pb on both sides of the transposon ends and ii) the number of events. \*\*, between parentheses are indicated the number of events.

In essence, these results lead us to conclude that MER75 may be a more suitable *pble* DNA substrate for initiating integration events by Mm523, compared to MER75B, MER85, Looper, and *Tcr-pble*. Indeed, MER75 appears to be a DNA substrate analogous to *Ifp2* for integration. However, it is crucial to note that integration assay results indicated that MER75, MER75B, and MER85 all exhibited similar qualities as *pble* DNA substrates for active integration by PGBD5 isoforms, facilitated by transposase-dependent integration.

These findings also shed light on the intriguing question of how PGBD5 can mobilize *pbles* such as MER75 or MER85. Notably, MER85 and MER75 share a common *pble* origin with PGBD3 and PGBD4, respectively (for an in-depth review, see supplementary data 2 in (2)). PGBD3 and PGBD4 genes are constitutively transcribed into mRNA and translated into protein in various sources of HeLa cells (for PGBD3, refer to https://www.proteomicsdb.org/protein/71758/expression and https://www.proteinatlas.org/ENSG00000225830-ERCC6/cell+line, and for PGBD4, see https://gpmdb.thegpm.org/∼/dblist_label/label=ENSP00000380872&proex=-1 and https://www.proteinatlas.org/ENSG00000182405-PGBD4/cell+line). Certain laboratory strains of HeLa cells were also found to transcribe the *PGBD5* gene and translate it into PGBD5 isoforms (https://gpmdb.thegpm.org/∼/dblist_label/label=ENSP00000375733&proex=-1). These observations raise questions about the possibility that a transiently transfected source of PGBD5 might need to interact with proteins like PGBD3 and PGBD4 to mobilize elements such as MER75 and MER85 during our integration assays.

To address this issue, we scrutinized gene expression profiles of our HeLa cells. RNA-seq experiments were conducted on cells subjected to various stressors (heat shock, geldanamycin or radicicol treatments) and compared to unstressed cells. Intriguingly, *PGBD4* was consistently transcribed irrespective of the treatment. *PGBD3* and *PGBD5* were not transcribed in unstressed cells, and no activation of their transcription was observed under stress conditions (supplementary data 3). These observations align with the hypothesis that an interaction between PGBD5 and other cellular factors may be necessary for mobilizing MER75 elements. However, the absence of detectable PGBD3 transcripts in HeLa cells suggests that PGBD3 is not involved in the interaction between PGBD5 and MER85.

### PGBD5 binds to chromosomal human *pbles* in certain human cells

To investigate the binding of PGBD5 to specific chromosomal loci in cells such as those containing human *pbles* or other discrete sequences, we undertook on a series of ChIP-seq experiments. Regrettably, due to the commercial unavailability of specific anti-PGBD5 antibodies, we sought to produce PGBD5-specific immune sera via DNA vaccination technology, ICANtibodies^TM^, a strategy we had successfully employed in a prior study involving another human domesticated transposase, SETMAR (19). The DNA vaccination was performed using pVAX plasmid encoding mouse Mm409 which exhibited a protein sequence identical to all other potential isoforms. Therefore, it was anticipated that our anti-PGBD5 sera would contain antibodies capable of binding to all other potential isoforms. Our initial attention focused on verifying the presence of anti-PGBD5 and non-specific antibodies. This verification process commenced with Western blot analyses (Figure 5A), employing protein extracts sourced from HeLa cells transfected with a 5Xmyc-Mm523 fusion construct.

**Figure 5.**
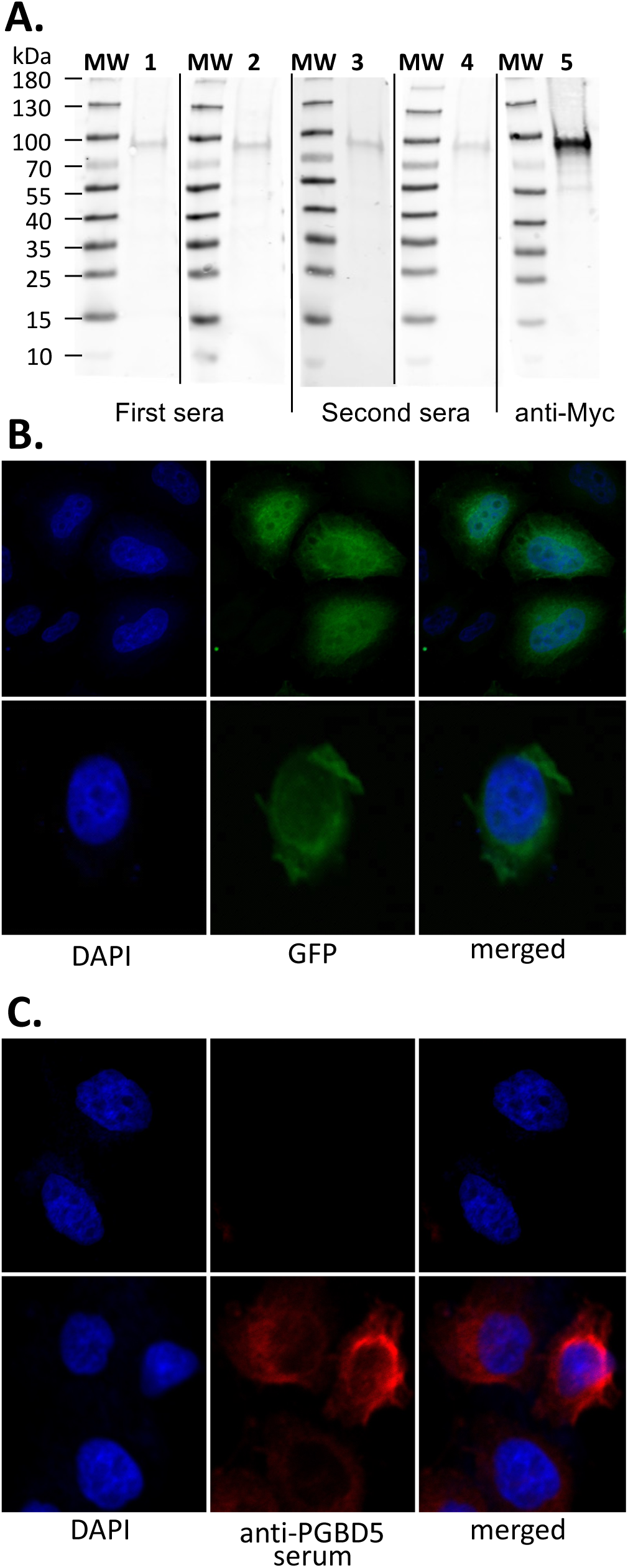
Properties of anti-PGBD5 pA in sera of SWISS mice. (**A**) Presence of pA directed against linear PGBD5 antigens. The Western blot analyses were done using protein extract from HeLa cells that were transfected 48 hours earlier with a pCS2 plasmid expressing 5Xmyc-Mm523 PGBD5 isoform. Lanes 1 and 3 were assayed sera harvested on day 21, lanes 2 and 4 on day 56. Sera were used in a dilution of 1/250. Lane 5 is shown a positive control using an anti-c-myc monoclonal antibody. MW, molecular weights ladder. Following this result, SW3 and SW4 sera were gathered for further validation in confocal microscopy. Preimmune sera of the two mice were also gathered and used as a control. (**B**) Cellular localization by confocal microscopy of Mm409-GFP in transiently transfected into HeLa cells. Top panel of three pictures: HeLa cells that were transfected 48 hours earlier with a pCS2 plasmid expressing a GFP plasmid. Bottom panel of three pictures: HeLa cells that were transfected 48 hours earlier with a pCS2 plasmid expressing the Mm409-GFP fusion. (**C**) Cellular localization by confocal microscopy of Mm409-GFP in transiently transfected into HeLa cells. Top panel: HeLa cells that were transfected 48 hours earlier with a pCS2 plasmid expressing the Mm409 PGBD5 isoform and revealed with the pre-immune serum. Bottom panel: HeLa cells that were transfected 48 hours earlier with a pCS2 plasmid expressing Mm409. The presence of PGBD5 is revealed with our anti-PGBD5 serum. In (**B**) and (**C**), the left pictures show the nuclear genomic DNA staining by DAPI, the middle picture show the GFP (top) or CY3 (bottom) fluorescence, the right pictures correspond to merged images.

These initial analyses revealed that two anti-PGBD5 sera contained pA antibodies directed against linear PGBD5 epitopes. Alas, these antibodies proved to be present in insufficient concentrations or had weak avidity towards their linear antigens. This deficiency rendered them inadequate for the detection of PGBD5 in samples featuring low PGBD5 concentrations, such as tissue biopsies expressing endogenous PGBD5. Consequently, we gathered these two sera and we verified the presence of pA antibodies directed against 3D antigens in the resultant composite serum using a multi-pronged validation approach. At first, we confirmed the presence of anti-PGBD5 pA directed against 3D antigens by comparing the profiles of cellular localization obtained with cells affixed to slides following transfection of a source of PGBD5-GFP or a source of PGBD5 revealed with our serum. HeLa cells were therefore transfected on one hand with pCS2-Mm409-GFP (Figure 5B and S2A) or pCS2-Mm409 (Figure 5C and S2B) and, on the other hand, with pCS2-Mm523-GFP (Figure S2C) or pCS2-Mm523 (Figure S2D and S2E). The outcomes conclusively established that our serum contained a sufficient concentration of pA directed against 3D PGBD5 antigens, facilitating the detection of PGBD5 within transfected HeLa cells. In addition, the profiles of PGBD5-GFP and PGBD5 revealed with our serum were similar, indicating that pA directed against 3D PGBD5 antigens were specific. Notably, our results unveiled that PGBD5 isoforms were distributed within the cytoplasm and nucleus. Intriguingly, the nuclear localization of both PGBD5 isoforms deviated from the typical patterns observed with most eukaryotic transposases (10,37). Both isoforms exhibited a preferential localization near the nuclear membrane, with a non-uniform dispersion extending into the nucleoplasm.

To ascertain the sensitivity of our serum, we analyzed a panel of cell lines. Specifically, we assessed cells lacking endogenous PGBD5 expression, such as HeLa and HEK293T cells, and cell lines featuring low transcription rates, including T98G, H4, 8-MG-BA, and HCT116. Our comprehensive assessment yielded no discernible background signals or spurious signals (Figure 5C, S3A-C) in our negative controls, encompassing HeLa and HEK293T cells, as well as the pre-immune sera. Furthermore, our serum reliably confirmed the presence of PGBD5 in both the cytoplasm and the nucleus of T98G, H4, and 8-MG-BA cells (Figure S3A-C). Significantly, the signal within the nuclei exhibited non-homogeneous distribution profiles. In the case of HCT116 cells, the signal detected following staining with our anti-PGBD5 serum was weak but significantly stronger than the signal obtained with the pre-immune serum. Nonetheless, an analysis of transcriptomic datasets from the BioProject PRJNA296592 (https://www.proteinatlas.org/ENSG00000177614-PGBD5/cell+line) indicated that *PGBD5* was transcribed and translated in the HCT116 cell line (https://gpmdb.thegpm.org/∼/dblist_label/label=ENSP00000375733&proex=-1), albeit likely at a lower level as compared to T98G, H4, and 8-MG-BA cells.

Our ChIP-seq analyses encompassed HeLa, HEK293, T98G, H4, 8-MG-BA, and HCT116 cell lines. Additionally, we revisited ChIP-seq data previously obtained with G401 cells (11) and subjected them to our computational pipeline. Statistically significant chromatin peaks for each cell line were documented in supplementary data 4. Remarkably, HeLa and HEK293-T cells, constituting our negative controls, yielded a mere four and seven peaks, respectively. H4 and 8-MG-BA cells failed to yield any significant chromatin peaks. Considering our findings in cells stained with our anti-PGBD5 serum (Figure S3B and S3C), this implied that PGBD5 exhibited limited affinity for chromatin or DNA within the nuclei of these two cell lines and/or the generated immune serum has insufficient affinity to chromatin-bound PGBD5.

In striking contrast, 283, 618, and 1306 significant chromatin peaks interspersed among chromosomes were obtained in the three remaining cell lines, T98G, G401, and HCT116, respectively. Notably, our analyses of G401 cells (11) concurred with our findings, as we obtained 100% congruence with the 618 peaks using PePr, although our results identified eight paired peaks, subsequently merged into four peaks compared to the earlier dataset. Importantly, no overlap was identified among the peaks obtained across these three cell lines.

We then investigated the presence of conserved sequence motifs within PGBD5-bound chromatin regions. At first, this was pursued separately for each cell line, employing RSAT and the MEME-suite. Astonishingly, only one remarkably significant and recurrent motif was detected in 87 T98G and 77 G401 peaks. No such motif was detected in HCT116 cells. This (TGGAA)n motif recurred at high frequency, with 5734 and 8302 occurrences in T98G and G401 peaks, respectively (Figure 6). This motif bore a striking resemblance to the well-documented tandemly repeated motif that constitutes the core of human centromeric satellite DNA III, where it serves as the principal component (41–43).

**Figure 6.**
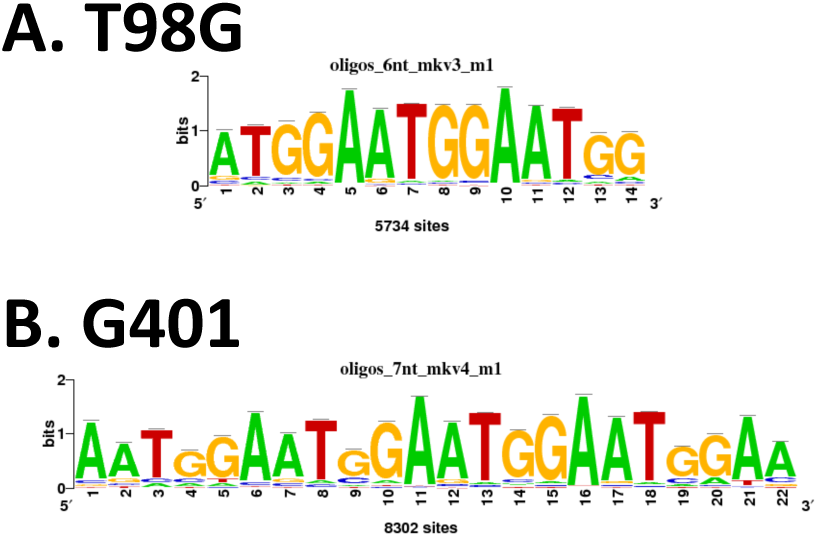
Motifs detected with RSAT and MEME in sequences overlapping ChIP-seq peaks in T98G and G401 cells. Main repeated motifs found in T98G (**A**) and G401 (**B**) colocalizing with ChIP-seq peaks. Their number of occurrences among sequences are indicated below each logo graphs. These motifs corresponded to the main motif consisting of the human centromeric satellite DNA III (41–43).

The implications of this discovery are twofold. Firstly, it’s worth noting that human centromeric satellite DNA III is specifically organized within the nuclear lamina that lines the inner facet of the nuclear membrane. Secondly, these recurrent motifs are recognized for their ability to assemble intrastrand dyadic DNA structures, including hairpins and cruciform DNA configurations. As thereafter confirmed by the other results, this second characteristic may likely be more important for the physical recruitment of PGBD5 than the presence of the tandemly repeated TGGAA motif.

To ascertain whether other repeats consistently overlapped with ChIP-seq peaks, we conducted additional analyses utilizing hg38 repeat annotations. While certain overlaps with abundant repeats in the human genome, such as AluI and L1 elements, did emerge, permutation tests underscored their lack of statistical significance (p>0.5). Significantly, this analysis consistently revealed four categories of repeats that prominently overlapped with PGBD5 ChIP-seq peaks (Figure 7): simple sequence repeats (SSR, encompassing mini and microsatellites), and three types of human *pble*, MER75, MER85, and Looper. Intriguingly, no overlap was identified with MER75B. The human *pble* annotation in the RepeatMasker (RM) annotation was therefore improved then verified (supplementary data 5) in order to improve the precision of these observations.

**Figure 7.**
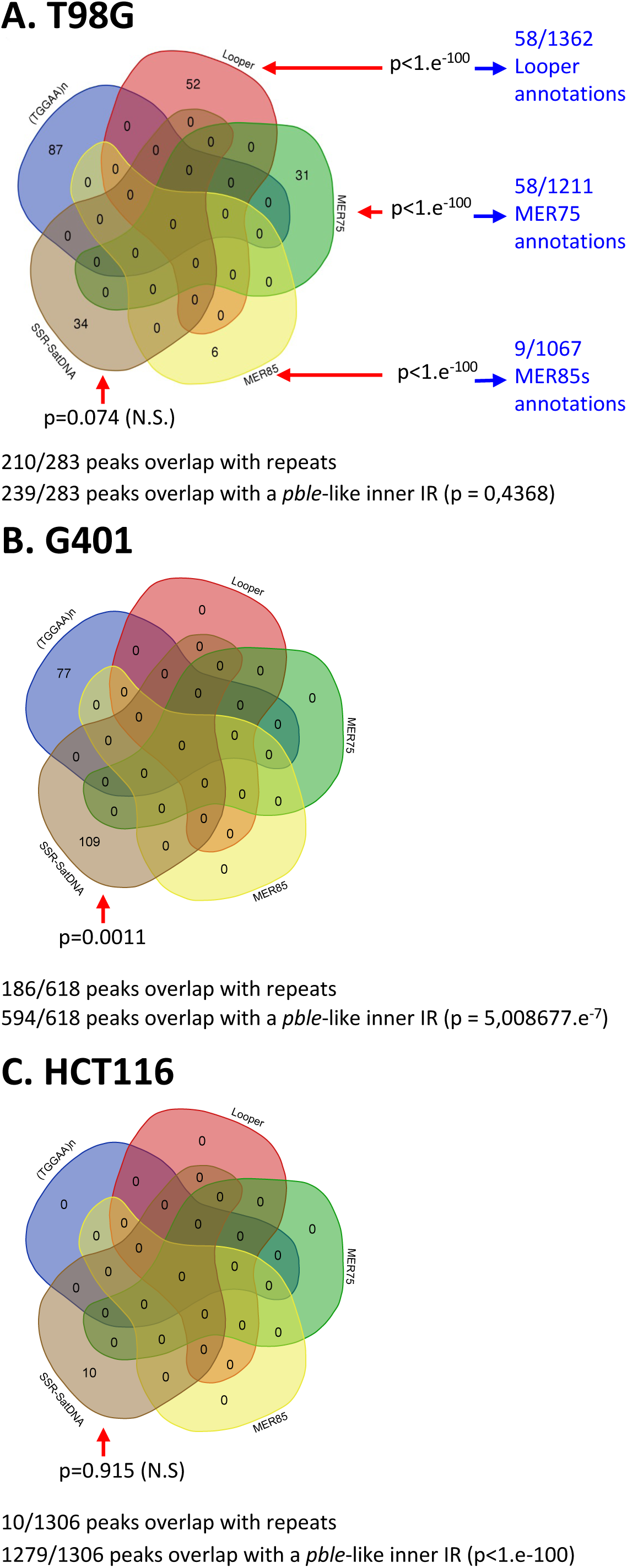
Repeat annotation of sequences overlapped by ChIP-seq peaks in T98G (A), G401 (B) and HCT116 (C) cells. The annotation results are represented in Venn diagrams with five leaves: MER75 (green area), MER 85 (yellow area), Looper (red area), (TGGAA)_n_ repeats (blue area), SSR (Simple sequence repeats; brown area). The numbers of co-occurrences are indicated within areas. Red arrows and p values indicated the probability to obtain co-occurrences by chance. On the right hand are indicated in blue, the numbers of co-occurrences with respect to the number of *pble* annotations (supplementary data 4). Below each Venn graph is indicated the number of peaks overlapping with one of the five repeats and the number of peaks overlapping with an IR displaying sequence features of a *pble*-like inner IR. Beside is indicated the probability to obtain this result per chance.

These studies identified PGBD5 chromatin regions that overlapped with SSRs (Figure 7: 34, 109 and 10) across all three cell lines. SSRs are recognized for their propensity to create intrastrand dyadic DNA structures, such as hairpin and cruciform DNA configurations (44). In contrast, PGBD5 peaks overlapping with MER75, MER85, and Looper elements were exclusive to T98G cells (Figure 7A). MER75, MER85, and Looper elements (Figure 8) share a common characteristic with invertebrate *pbles*, namely, the presence of STIR or IIR at both their ends (2). It is worth noting that MER75B exhibits subterminal inverted repeats (IRs) only at one of its subterminal regions (Figure 8). Under conditions that favor the assembly of non-B DNA configurations (45,46), these STIR and IIR could theoretically facilitate the formation of intrastrand dyadic DNA structures, such as hairpin and cruciform DNA configurations. Permutation tests were employed to assess whether the overlaps between peaks and these three types of repeats carried statistical significance. In T98G, overlaps with MER75, MER85, and Looper were found to be highly significant (Figure 7), but this significance was not observed in the case of SSRs. For G401, overlaps with SSRs were indeed significant, but not in HCT116.

**Figure 8.**
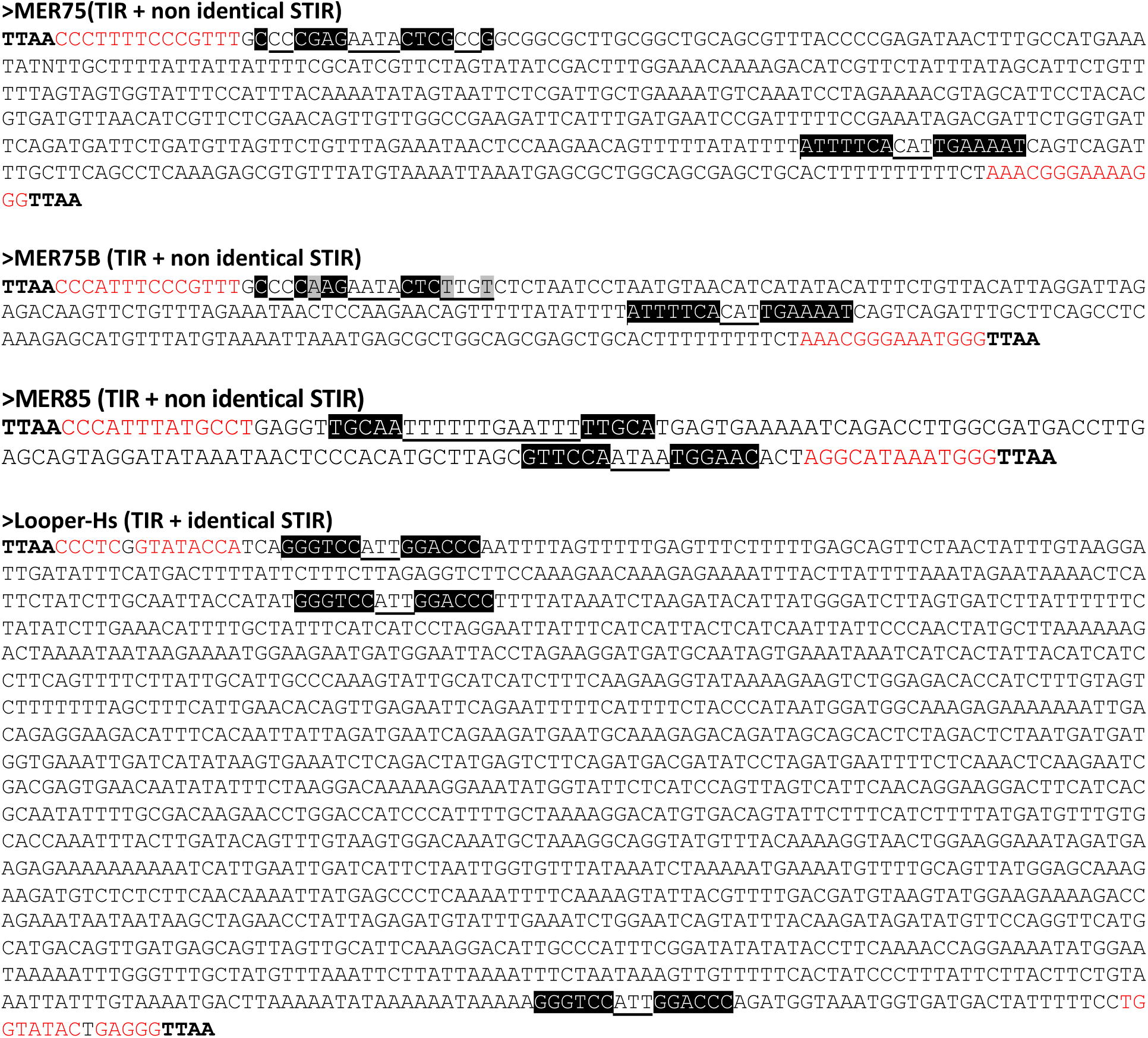
Sequence features of human *pble*s, MER75, MER75B, MER85 and Looper. The motifs corresponding to the target site duplication of *pble*s, TTAA, are in bold, the inverted terminal repeats are typed in red, and the inner IRs in subterminal regions are underlined and nucleotides within palindromic repeats are typed in white and highlighted in black. In the MER75B sequence are highlighted in grey divergent nucleotides that switch off the palindromic organization of the inner IRs in subterminal regions present at the 5’ end of MER75.

Recognizing that the common feature among the five kinds of repeats ((TGGAA)n, SSR, MER75, MER85, and Looper) was their ability to facilitate the formation of intra-strand dyadic DNA structures, we proceeded to annotate inverted repeats (IRs) within hg38. This annotation aimed to determine whether at least some of the sequences that did not overlap with these five repeats in T98G, G401, and HCT116 might overlap with one or more IRs. The parameters used for this analysis were consistent with those exhibited by *pble* STIR and IIR motifs (2). These parameters encompassed a repeat size ranging from 5 to 15 nucleotides, a spacer between pairs of IRs spanning from 2 to 10 nucleotides, and a number of mismatches between both repeats, ranging from 0 (for repeats of at least 5 nucleotides or more) to 1 (for repeats of at least 6 nucleotides or more). Our analysis revealed that a substantial proportion of peaks, specifically, 85%, 96%, and 98% in T98G, G401, and HCT116, respectively, overlapped with *pble*-like IRs (Figure 7). Importantly, these results garnered robust statistical support through permutation tests conducted in G401 and HCT116 cells. However, the statistical significance in T98G hovered below the threshold. This result can be attributed to the higher prevalence of peaks overlapping with (TGGAA)n repeats in T98G (comprising 28% of peaks), as compared to G401 (12%) and HCT116 (0%). Notably, the potential for IRs to facilitate the formation of dyadic structures was not considered in the annotation procedure for satellite DNA III that we employed.

Finally, we sought to investigate whether the distribution of peak features and read density in T98G and G901 was contingent upon the type of repeat they intersected. This analysis uncovered several notable trends. For instance, peaks that intersected with SSRs and satellite DNA III motifs exhibited significantly greater widths (Figure 9A and C; with a median around ∼2 kbp in T98G and G401) compared to those overlapping with *pbles* in T98G (Figure 9A, with a median ranging from 600 to 800 bp), and noTR peaks in G401 (Figure 9C, with a median around ∼500 bp). This was expected since SSRs and satellite DNA can span long stretches of tandemly repeated motifs, offering multiple binding sites for PGBD5. Similarly, since MER75, MER85, and Looper elements can potentially harbor binding sites at both ends, depending on their genomic context, it is unsurprising that the peaks overlapping these elements exhibited widths close to their size.

**Figure 9.**
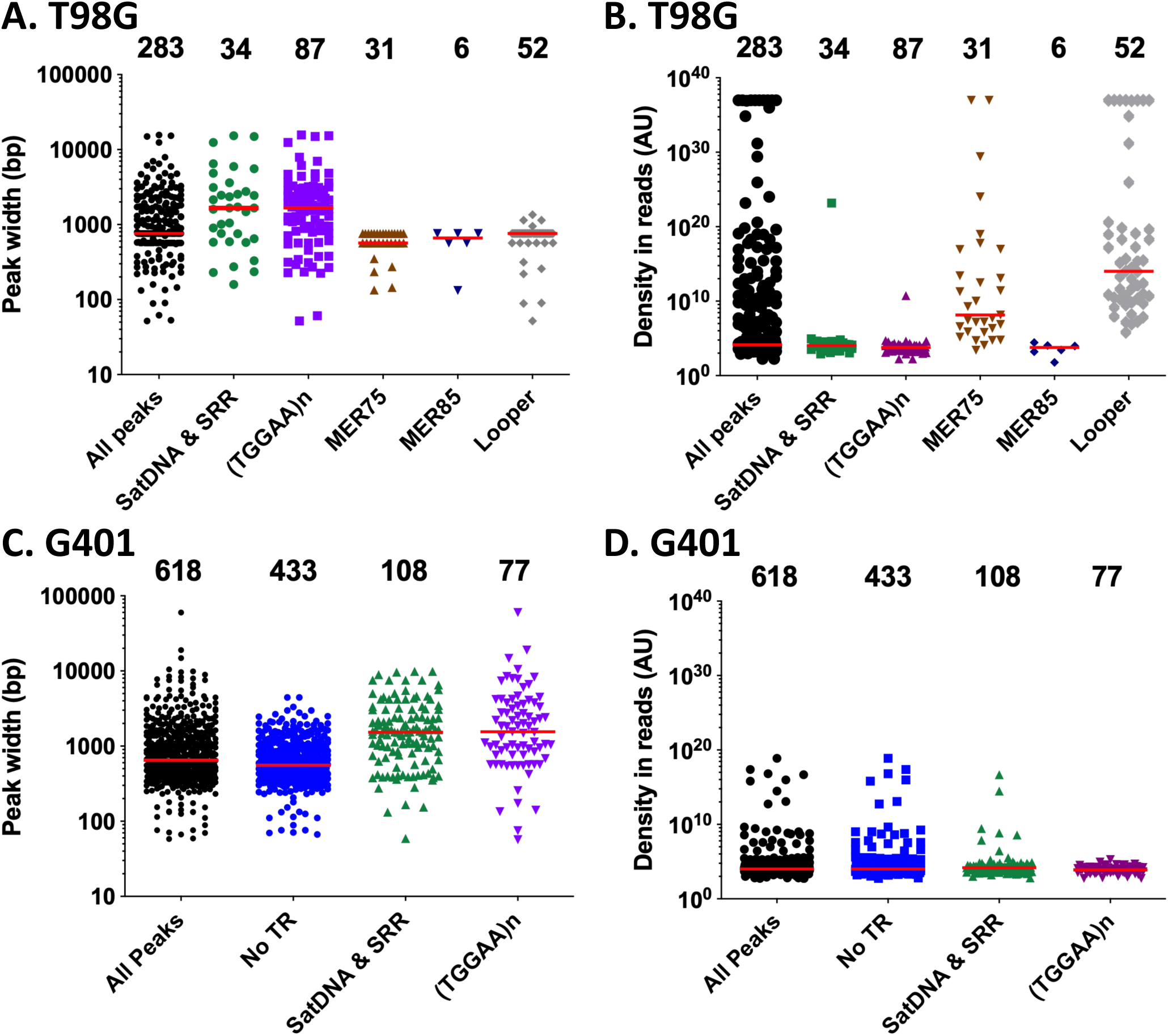
Graphic representation of peak widths (A and C) and read densities (B and D) between the different categories of repeats overlapped by peaks in T98G and G904 cells. In (**B**) and (**D**), the density of each peak was calculated from the number of reads divided by its peak width (in nucleotides). Results were expressed in arbitrary units. No TR (tandem repeats) corresponded to peaks not overlapping (TGGAA)_n_, SatDNA (satellite DNA) and SSR sequences. The numbers of peaks for each category were indicated on the top of each graphic. The red lines represented the median.

Furthermore, we observed that MER75, and more prominently, Looper, displayed higher median read densities compared to MER85 and other categories of tandem repeats in T98G cells (see Figure 9B). In contrast, in G401, the density of reads across all peak categories exhibited a more uniform distribution (Figure 9D). It is worth acknowledging that while these differences may partly stem from PCR biases during Illumina sequencing library preparation, they align with the hypothesis that MER75 and Looper sequences represent suitable DNA substrates for PGBD5. However, this could also be explained by a “site occupancy” hypothesis in which MER75 and Lopper loci would be more accessible in the chromatin of a cell population than those containing MER85, (TGGAA)n and SatDNA and SSR repeats. Consequently, they may be more frequently bound by PGBD5 in T98G cells.

## Discussion

This work represents the first demonstration that human PGBD5 can interact with human *pbles*, facilitating several alternative forms of genomic transposon integration in experimental integration assays. Furthermore, we have established the ability of PGBD5 to bind to chromatin at loci harboring human *pble* copies in human T98G cells, as well as at sites characterized by the presence of tandemly repeated motifs such as satellite DNA III or certain SSRs in both T98G and G401 cells. Our data also underscore a robust correlation between PGBD5’s binding to chromosomal loci and the presence of *pble*-like inner IR within the sequences of bound loci. Interestingly, this correlation extends to loci devoid of SSRs, satellite DNA III, and human *pble* sequences.

In comparison to a conventional *pble* transposase like the insect PB, the activities of human PGBD5 appear more intricate. Firstly, we currently lack a system suitable for studying its specific biochemical activities via *in vitro* assays. Secondly, certain facets of PGBD5 activities may be residual, persisting despite its 500 million years of evolution. Evidently, its amino acid sequence exhibits exceptional conservation among mammals, with over 93% identity within the last 409 C-terminal residues. This high degree of conservation underscores a potent selection pressure that can now be attributed to its pivotal role(s) in host development and physiology (47). This conservation is likely associated with its integration activity of *pble* DNA fragments, which we have found to be at least 10 times weaker than that of insect PB with the active mammalian PGBD5 isoforms, Mm409 and Mm523. Consequently, this prompts questions regarding PGBD5’s evolved activities within its vertebrate hosts. Is it a site-specific transposase, does it function in alternative forms of DNA rearrangements via its cleavage activities, or involve as of yet more complex chromatin mechanisms? Might it resemble an excisase akin to RAG1/2 (9), utilizing its own DNA targets for deletional recombination, or alternatively, could it function as another type of endonuclease, or has it evolved into a protein with its primary domain, previously known for its Ruvc/RNAseH fold, having lost its classical DD[E/D] catalytic triad and acquired a divergent one during its very early evolutionary divergence (12)? It must be noted at this point that the unique catalytic triad of PGBD5 is not exclusive to this protein, as at least two other domesticated eukaryotic transposases exhibit DNA binding, DNA cleavage, and/or DNA strand transfer activities with non-canonical catalytic triads. The first occurs in SETMAR, which is no longer fully functional due to the substitution of one of the essential aspartates with an asparagine. This substitution is likely not a simple D→ N replacement, but rather the result of an evolution involving the initial loss of the catalytic D, followed by partial structural recovery through the emergence of a non-catalytic N residue. This suggests that SETMAR underwent a more complex evolutionary trajectory, possibly reflecting selective pressure to retain a vestigial transposase fold without restoring full catalytic function (48,49). SETMAR has also been shown to mediate SETMAR-dependent plasmid integration in assays similar to those used with PGBD5 (50). The second set of proteins belongs to the Fanzor nucleases, derived from TnpB transposases (51). After the evolutionary loss of the second D residue in the DDD triad in two different Fanzor lineages, their evolution was likely marked by functional adaptations that enabled the restoration of another DED triad through the recruitment of two alternative E residues located near the lost D residue. Together, these examples highlight that the presence of a non-canonical catalytic triad within the sequence of a domesticated transposase can result from the more or less precise restoration of activities of its transposase ancestor. Lastly, it is also possible that PGBD5 recruits another nuclease to mediate its cellular genomic remodeling activities.

After millennia of evolution within vertebrate genomes, PGBD5 has also likely undergone co-evolution with its cellular environment. It may have acquired specific regulatory mechanisms governing its transcription, translation, conformational folding, post-translational modifications, and subcellular localization. Our results provide insights into this, showing that PGBD5 does not randomly localize and disperse within nuclei in transfected cells and cell lines. Moreover, our ChIP-seq findings suggest that PGBD5 can exist within nuclei without binding to DNA or chromatin in cell lines such as H4 and 8-MG-BA, indicating a potentially inactive configuration or state with non-chromatin functions. Lastly, considering that several PGBDs (also known as *piggyMac*-like proteins) collaborate in certain ciliate species to execute programmed DNA rearrangements, excising approximately 40,000 discrete DNA fragments during macronuclei maturation, another layer of complexity in the functionality of PGBD5 could involve its potential to interact with one or several of the four other PGBDs and/or host factors associated with DNA repair or the recognition of transcriptional units, as previously observed for PB (52,53).

### Integration by transposition, recombination, or PGBD5-dependent integration

Integration of DNA fragments carrying a selection cassette can occur through various mechanisms, including canonical ‘cut-and-paste’ transposition, recombination, or DNA breakage followed by repair. In the case of MER75, the number of integration breakpoints matching with a canonical transposition signature defined by precise target site duplications was more elevated than per chance. This finding suggests a physical interaction between PGBD5 and MER75, capable of facilitating proper transposition events, akin to our previous observations with *Ifp2* and *Tcr-pble* (2,10,11,12).

However, when we examined MER75B, MER85, and Looper, the results obtained from HeLa cells did not support the same conclusion. Nevertheless, we observed that PGBD5 increased their genomic integration through PGBD5-dependent integration events. Notably, this augmentation was only evident when *pble* ends were present, indicating that PGBD5 possesses the ability to interact directly or indirectly with MER75B, MER85, and Looper. Our ChIP-seq results in T98G cells further validate this interaction, particularly in the case of the last two *pble* elements. Collectively, these findings affirm that PGBD5 exhibits the capability to interact with *pble* elements, albeit with potentially varying affinities for each. Importantly, we also discovered that this property is not exclusive to PGBD5 but is likely shared by most *pble* transposases. Indeed, our results corroborate those obtained with *NlPLE25* (41), supporting that PB can interact with *pble* elements other than *Ifp2* and integrate them into chromosomes. This observation challenges the long-standing dogma in the transposon vector field, which postulated that a transposase of one transposon “species” would be restricted in its interactions with its conspecific elements.

It is worth noting that insect PB and human PGBD5 are not the first eukaryotic transposases to demonstrate the ability to interact with non-conspecific elements. This property has previously been shown in some *Tc1-mariner* elements that display relaxed sequence specificity (54). This property was even employed as a tool to discover new active transposases or to characterize the activity of domesticated transposases in the *hAT* and *P* transposon families (55–59). Therefore, our findings represent yet another instance of the flexibility inherent in a third family of eukaryotic transposons, the *piggyBac* family.

Moving beyond the notion of strict specificity in transposition, it is also important to reconsider the idea that transposases such as PB and Sleeping Beauty (SB) exclusively catalyze proper transposition events, avoiding so-called illegitimate integration. To embark on this exploration of PGBD5’s integration activities, it becomes essential to mine the existing literature in order to gauge the robustness of our understanding of what constitutes a canonical transposition event. This undertaking requires delving into two fundamental questions: i) is there any evidence to suggest that eukaryotic transposases might give rise to illegitimate integration during integration assays? and ii) are the methodologies currently employed in the literature adequate for the detection of non-canonical integrations?

To address the first question, we found a solitary published experimental result utilizing an assay tailored for detecting illegitimate integration events during the use of SB10 as a transposase source (Figure 4 in (60)). This assay involved a transposon donor plasmid carrying a NeoR cassette flanked by *Sleeping Beauty* ends, with an additional cassette in its backbone enabling the systemic expression of the thymidine kinase gene of herpes simplex virus type 1 (HSVTK). The utility of such a plasmid lies in its ability to selectively identify G418-resistant cells that have integrated NeoR into their chromosomes through proper *Sleeping Beauty* transposition or recombination. It also allows for the elimination of clones containing both NeoR and HSVTK, which would result from co-integration events, by monitoring G418 selection in the presence or absence of ganciclovir, which is toxic to cells expressing HSVTK. This assay was conducted across six different human and murine cell lines (60), and the results revealed that, depending on the cell line, between 10% and 85% of NeoR clones contained co-integrated NeoR and HSVTK sequences within their chromosomes. These clones arose from either the sole integration of the transposon donor plasmid through recombination or concurrent events of transposition and integration of the transposon donor plasmid through recombination and/or alternative modes of DNA rearrangements. Control experiments conducted in the same study (60) indicated that only 5% of NeoR clones resulted from the random integration of the transposon donor plasmid. Consequently, these results strongly support the idea that, depending on the cell line, 5% to 80% of NeoR clones integrated HSVTK via transposase-dependent integration events. To date, there have been no replications of these experiments involving *Sleeping Beauty* or other transposon vectors. Therefore, these results remain the sole evidence suggesting that, in an integration assay, a DNA cassette flanked by transposon ends can integrate at varying frequencies through canonical transposition, recombination, or transposase-dependent integration events, contingent on the cell type used, and likely independent of the used transposon vector.

In conclusion, several published studies shed light on the diverse mechanisms of integration and the flexibility exhibited by eukaryotic transposases, challenging long-held assumptions in the transposition field. Furthermore, it underscores the need for a comprehensive reevaluation of what constitutes a canonical or proper transposition event, as well as the methodologies employed to detect illegitimate integration. The initial investigations aimed at delineating the insertion profiles of *Ifp2* and *Sleeping Beauty* transposons commenced in the early 2000s. These pioneering studies employed methodologies such as inverse PCR coupled with sequencing, PCR with splinkerettes followed by NGS sequencing, or plasmid rescue combined with Sanger sequencing. When examining the PB/*Ifp2* vectors, these endeavors unveiled a landscape where a staggering 96-98% of *Ifp2* insertions into chromosomes resulted from canonical transposition. In this context, canonical transposition signifies integration without lesions occurring within *Ifp2* vectors and their termini. Furthermore, these integrations occurred precisely at a TTAA tetranucleotide, serving as the target site duplication (TSD). Importantly, there were no traces of the flanking plasmid backbone from the transposon-donor plasmid observed at the outer extremities of the TSD (e.g., references (61–65)). Remarkably, no instances of non-canonical integration were reported, a stark contrast to numerous studies demonstrating such events in their control experiments.

However, our recent findings have challenged this idea. We employed techniques like LAM-PCR and FLEA-PCR, utilizing sequence alignment and analysis algorithms optimized for the detection of structural variants and DNA breakpoints. In our investigations, we characterized 7623 insertion breakpoints (10): of these, 6828 (90%) resulted from canonical transposition at TTAA TSD, 543 (7%) occurred within divergent TSD tetranucleotides but were still the outcome of proper transposition events, as convincingly demonstrated (65). Intriguingly, 252 (3%) were situated within the transposon ends (187; 2%) or within the flanking plasmid backbone of the transposon-donor plasmid at the outer edges of the TSD (65; 1%). The discrepancy between these results and those obtained from earlier plasmid rescue and splinkerette methods can be attributed to the fact that the former techniques were primarily designed to establish the insertion profiles through canonical transposition, rather than to gauge all integration events. Non-canonical events were effectively filtered out through three distinct means: i) the transposon-donor plasmid was engineered to feature restriction sites at both outer edges of the TSD, facilitating the elimination of transposon ends still linked to the plasmid backbone through digestion in selected clones; ii) sequencing primers were employed with a 3’ end in very close proximity to the TIR ends; iii) during computational analyses, filtering steps were applied to eliminate non-transposition events. These considerations are essential when scrutinizing recent strategies not explicitly designed for assessing insertion quality (66) and when they are used extensively to evaluate the insertion profiles of novel vectors and transposases (67).

In terms of the precision of canonical transposition, previous studies mainly concentrated on repairing the transposon donor site post excision. However, one study (68) reported the occurrence of lesions at the transposon ends in the case of a non-*pble* DNA transposon, the *mariner mos-1* element. These lesions resulted from cleavage events executed by the MOS1 transposase at the ends of a *mos-1* element, known as *peach*, which was already integrated into a chromosome. Importantly, these cleavages were not resolved through the non-homologous end-joining (NHEJ) DNA repair pathway but rather through the homologous recombination (HR) pathway. HR (69) is a DNA metabolic process that facilitates high-fidelity, template-dependent repair or tolerance of complex forms of DNA damage, including DNA gaps, DNA double-stranded breaks (DSBs), and DNA interstrand crosslinks (ICLs). In cells, the activation of these two pathways depends on the cell cycle phase (70). This pattern of lesions at the ends of DNA transposons mirrors what has been observed with copies of two other *mariner* element “species” in the human genome, *Hsmar1* (see profile at https://www.dfam.org/family/DF0000212/seed and (19)) and *Hsmar2* (see profile at https://www.dfam.org/family/DF0000213/seed). In the broader context, DNA transposon copies with inner deleted segments are relatively rare in the human genome, most of them rather displaying with more or less extensively trimmed ends. Molecular processes underlying this phenomenon remain poorly understood, even though they align with what was observed with *mos-1*.

In the presence of PB, similar events may occur at the ends of *Ifp2* vectors. Following PB cleavage, HR could facilitate DNA repair, potentially utilizing as templates other molecules of transposon donor plasmid that were co-transfected in the cell nucleus. This could elucidate how deletions within the transposon terminal regions or the insertion of plasmid backbones can occur with the PB/*Ifp2* couple, as well as with the PGBD5/*pbles* pair. In addition, certain properties of insect PB and human PGBD5 might explain why the rate of illegitimate events is significantly higher with PGBD5 than PB. Our results indicate (Figure S1) that the rates of DNA cleavage by PB and Mm523 were quite similar under the experimental conditions of our integration assays. However, we also observed that the clonal capacity of human cells expressing Mm523 PGBD5 is approximately 12-fold lower than that of insect PB (2,10,17). Like SB (71), PB in the cell nucleus is physically associated with several key proteins involved in the NHEJ pathway (Ku70, Ku80, DNA-pKcs) (52). In accordance with this, a candidate B-Ku binding motif (72) is present on the PB surface in the 6X67 PDB structure, spanning residues 211 to 234 (<u>M</u>S<u>R</u>D<u>R</u>FD<u>FL</u>IR<u>C</u>L<u>R</u>MDDKSI, with conserved residues underlined). A similar motif is conserved in the same location in most *pble* transposases, but no such motif is found in PGBD5. Consequently, if PGBD5 interacts with the NHEJ machinery more loosely than PB, this could lead to more frequent repair of excision and insertion events mediated by HR, resulting in a higher occurrence of illegitimate integration events. Further investigations into PGBD5 and its interactions with DNA repair pathways will be necessary to gain a deeper understanding of its cleavage and strand transfer activities, as well as its relationships with repair mechanisms in the host cell machinery.

### PGBD5 displays complex genomic remodeling activities in human cells

As previously highlighted, our ChIP-seq findings across various cell lines suggest that the regulation of PGBD5 activities likely operates at multiple levels. Notably, we have observed instances where PGBD5 is present within nuclei without apparent binding to chromatin or DNA, as seen in H4 and 8-MG-BA cells. Initially, we interpreted this as the existence of two post-transcriptional isoforms or structural conformers of PGBD5, with one capable of binding to chromatin or DNA and the other not. However, we must now consider two additional non-exclusive and complementary alternatives.

The first possibility is that PGBD5 could bind to *pble*-like IRs that are widespread within chromosomes, as well as to STIRs or IIRs within *pble* sequences. To delve deeper into this possibility, we conducted electrophoretic mobility shift assays using nuclear extracts from HeLa cells transiently transfected with each of the four 5Xmyc-PGBD5 isoforms and DNA probes corresponding to both ends of *Ifp2* and MER75. These conditions theoretically should have allowed us to detect shifted bands, whose specificity could subsequently be confirmed through supershift assays employing anti-myc antibodies as previously performed for another purpose (73). No shifted bands were observed under our experimental conditions,. It is important to note that the STIRs or IIRs in our DNA probes adopted a B-DNA configuration. Our analyses of sequences overlapped by ChIP-seq peaks in T98G, G401, and HCT116 cells have revealed that their common feature is not the presence of (TGGAA)n motifs, but rather sequences which have been described to form intrastrand dyadic DNA structures such as hairpins and cruciform DNA structures under specific constraints, such as DNA superhelicity (45,46). Given that *Ifp2* and MER75 contain STIRs or IIRs, it becomes imperative to assay PGBD5 binding using nucleic acid probes with ends that exhibit a stable fold mimicking hairpin and cruciform structures at the locations of their STIRs or IIRs. In theory, such probes could be synthesized using convertible and conformationally constrained nucleic acids (C2NAs) (74–77). To our knowledge, this has not been attempted with DNA fragments as long as those needed for these assays and would necessitate a substantial research effort involving specialists in oligonucleotide conformational synthesis. Nevertheless, confirming whether DNA configuration regulates PGBD5 binding is crucial, as it may provide insight into why sites bound in one cell line (such as some human *pbles* in T98G) are not bound in other cell lines (like G401 and HCT116).

The second alternative worth considering is that PGBD5 may interact with other cellular factors including other PGBD proteins to bind to its DNA targets. Currently available transcriptomic and proteomic data from databases indicate that PGBD1, PGBD2, and PGBD4 are expressed in cell lines such as HeLa, HEK293, T98G, H4, 8-MG-BA, HCT116, and G401, whereas PGBD3 is not. However, if these proteins share interaction properties with DNA or chromatin similar to and complementary with those of PGBD5, extensive mutant and complementation assays will be required to unravel their roles in integration and binding processes. Initially, it is imperative to ascertain whether they indeed interact. To address this, studies employing pull-down and ChIP-seq experiments will be necessary using appropriate antibodies, but they may become more intricate if PGBDs are found to exist in cells as both active and inactive isoforms or conformers.

Finally, we must also discuss PGBD5 chromatin binding sites identified in T98G, G401, and HCT116. If PGBD5 functions as a simple or complex nuclease, either on its own or with cellular cofactors, that can be in at least two different states, active and inactive, these binding loci must be considered under two non-exclusive hypotheses. At first, if PGBD5 is active when bound to chromatin, it should be expected to find 100% chromosomal regions overlapped by ChIP-seq peaks exhibiting nuclease-dependent rearrangements because of the ages of these cell lines (at least 45, 43 and 37 years-old for T98G (78), G401 (79) and HCT116 (80), respectively), and the many cell divisions they have undergone. This is not what we observe; Illumina reads are evenly distributed. This therefore supports that nuclease activities are not initiated when PGBD5 is bound to these loci. Consequently, it is possible that some or all of these binding sites may not be closely associated with recombination events but rather serve as storage sites where PGBD5 is held in reserve and not actively participating in recombination or involving other chromatin functions. This phenomenon has previously been observed in nuclei with transcription factors (81,82). The second hypothesis is that PGBD5 is bound to potent recombination sites as an inactive configuration due to post-translational and-or conformational modifications. Understanding how cells precisely control the availability of PGBD5 in its active and inactive forms within nuclei therefore represents an additional layer of biological complexity that must be explored in future studies.

### Consequences in translational transposon applications

“Cognate restriction of transposition” by DNA transposases is not a discrete phenomenon, but rather a relative activity dependent on transposase concentration, cellular factors, and DNA substrates. This is well established for many sequence-specific nucleases, such as restriction endonucleases for example, where they bind and cleave both higher affinity (specific) and lower-affinity sequences to varying degree, as determined by relative binding affinities, catalytic activities, and solution conditions. Likewise, overexpression of diverse DNA transposases has been extensively described causing DNA damage and cytotoxicity. The ectopic expression of PGBD5 can transform human cells due to the induction of oncogenic DNA rearrangements. A recent clinical trial of chimeric antigen receptor T (CART) cells engineered using *piggyBac* transposition and administered to patients reported two cases (20% of the patient cohort) of CART cell lymphomas (83,84). Resultant CART cell lymphomas exhibited high numbers of transposon copies, but no evidence of insertions into known oncogenes, nor changes in expression of transposon insertion-associated genes. In contrast, CART cell lymphomas had high genomic copy number changes, consistent with oncogenic genomic rearrangements, either from the genomic transposase activity (85), and/or another cellular mutational process such as those caused by variable efficiency of DNA repair (86). It is currently not possible to discriminate between these possibilities. While this malignant transformation is unlikely to be caused by PGBD5 which is not constitutively expressed in most lymphoid cells analysed to date, additional studies will be needed to definitively exclude this possibility in case PGBD5 is expressed inducibly upon inflammatory or developmental cues. Alternatively, in addition to canonical transposition, PB itself may also promote alternative forms of DNA rearrangements. Importantly, these results emphasize the continued need for critical evaluation of cellular activities of heterologously expressed DNA transposases (87,88). We hope that future collaborative research will continue to define the specific mechanisms and functions of these semantically simple, but biologically complex molecules.

## Data availability

All raw and processed data are available through the European Nucleotide Archive as BioProjects PRJEB36231 to PRJEB36234 for LAM-PCR datasets, PRJEB64412 for ChIP-seq datasets, and PRJEB64412 for RNA-seq. The data underlying this article for the analyses of the integration breakpoints, RNA-seq in HeLa cells and the ChIP-seq peaks are available in the article and in its online supplementary data 2, 3and 4. Annotations of human PBLEs in hg38 are available in supplementary data 5. Annotations of PBLE-like inner IRs in hg38 can be downloaded at https://doi.org/10.5281/zenodo.8281129.

## Supplementary data

Supplementary Data are available at NAR Online.

## Author contributions

Linda Beauclair : Conceptualization, Formal Analysis, Investigation, Data curation, Validation. Laura Helou : Conceptualization, Formal Analysis, Investigation, Software, Data curation, Validation. Florian Guilllou and Hughes Dardente : Formal Analysis, Investigation, Validation, Writing – review & editing. Thierry Lecomte : Funding acquisition, Writing – review & editing. Alex Kentsis : Writing – review & editing. Yves Bigot : Conceptualization, Formal Analysis, Methodology, Validation, Supervision, Writing – original draft.

## Acknowledgements

We thank members of our groups for critical feedback.

## Legends supplementary figures and data

**Figure S1. Impact of PGBD5 isoforms and PB variants on rates of H2A.X phosphorylation and survival in HeLa cells.** (**A**), normalized α-tubulin amount calculated from quantification on western blots made with protein extracts from 10^5^ HeLa cells transfected with 400 ng and a transposase/transposon donor ratio of 1 and, (**B**), graphic representation of H2A.X-P/α-tubulin. Transfection rate was ∼75% and was evaluated in parallel using a pGL3-DsRed-Kan (a gift of Pr Nicolas Mermod, Lausanne, Switzerland). (**C**) Western blot analyses of phosphorylated γ-H2AX, α-tubulin, and PRPF19 in HeLa cell extracts transfected with each of the four PGBD5 isoforms and the 3 PB variants. Each protein extract was prepared 24 hours post transfection from the complete cellular content of one well of a 24-well plate. In each plot, the red lines represented the median calculated from 5 experiments fully replicated. In A and B, data were normalized on pBSK and in C and D on the mock. * and brackets, and NS indicated a significant and a non-significant Krustal-Wallis test (p>0.05) between results obtained with PB, PB.1-558, PB.NLS-1-558, Mm523, Mm409, Hs455 or Hs554, and pBSK. In (**C**), for the revelation of phosphorylated γ-H2AX, α-tubulin and PRPF19, mouse anti-phospho-H2A.X and anti a-tubulin antibodies were used while and a rabbit polyclonal anti-PRPF19 was used. Secondary antibodies were a donkey anti-mouse-IR800 and a donkey anti-rabbit-IR700 (LICOR Biosciences, Lincoln, NE, USA). The picture corresponded to a fusion of those obtained with both dyes. The arrow in red located the α-tubulin, in green PRPF19 and in red the phosphorylated γ-H2AX. In (**A**) and (**B**), counts were done separately for both dyes. In all, these data indicated that i) the transfection affected the H2A.X-P/α-tubulin and the normalized α-tubulin amount (i.e. the cell survival and growth) since they were significantly different with those of control done with pBSK, ii) PB, PB.1-558, PB.NLS-1-558, Mm523, Mm409, Hs455 and Hs554 displayed a genotoxic activity that affected the cell survival and-or growth.

**Figure S2. Cellular localization by confocal microscopy of Mm409-and Mm523-GFP isoforms and their GFP fusions into transiently transfected HeLa cells.** (**A**), the two horizontal panels of three pictures were a complement of those shown in Figure 5B. They showed HeLa cells that were transfected 48 hours earlier with a pCS2 plasmid expressing the Mm409-GFP fusion. The left pictures showed the nuclear genomic DNA staining by DAPI, the middle pictures showed the GFP fluorescence signal that depicts the distribution of Mm409-GFP within the cell, the right pictures corresponded to mezrged images. Images showed that GFP-Mm409 signal was mainly located in the cytoplasm around the nuclear membrane, on the inner side of the nuclear membrane. The GFP-Mm409 signal was also present in the nucleoplasm. However, the signal was not diffuse and it appeared different from one nuclear region to another. (**B**), the two horizontal panels of three pictures were a complement of those shown in Figure 5C. They showed HeLa cells that were transfected 48 hours earlier with a pCS2 plasmid expressing Mm409. The left pictures showed the nuclear genomic DNA staining by DAPI, the middle pictures show the CY3 signal that depicted the distribution of Mm409 within the cell, the right pictures corresponded to merged images. Images showed that Mm409 is cytoplasmic and nuclear, mainly located around the nuclear membrane. The CY3 fluorescence signal also indicated that Mm409 was present in the nucleoplasm, the signal being not diffuse and appearing to be differently intense from one nuclear region to another and even have the shape of flame from the nuclear membrane toward the nucleoplasm centre. In all, the localization profiles A and B and in Figure 5B and C obtained in transiently transfected HeLa cells were similar. By comparison with the profiles of the PB (10) and *mariner* (37) transposases, the localization of PGBD5 was not exclusively nuclear, its distribution in nuclei being enriched onto the nuclear membrane and not diffuse within the nucleoplasm. (**C**) The three horizontal panels of three pictures showed HeLa cells that were transfected 48 hours earlier with a pCS2 plasmid expressing the Mm523-GFP fusion. The left pictures show the nuclear genomic DNA staining by DAPI, the middle pictures show the GFP fluorescence signal that depicts the distribution of Mm523-GFP within the cell, the right pictures correspond to merged images. Observations were similar to those shown in (**A**). (**D**) The three horizontal panels of three pictures show HeLa cells that were transfected 48 hours earlier with a pCS2 plasmid expressing Mm523 and revealed with our anti-PGBD5 serum. The left pictures show the nuclear genomic DNA staining by DAPI, the middle pictures show the CY3 signal that depicts the distribution of Mm523 within the cell, the right pictures correspond to merged images. Observations were similar to those shown in (**B**). (**E**) Z-stack imaging in 22 pictures (there are no pictures in 23 and 24) of HeLa cells that were transfected 48 hours earlier with a pCS2 plasmid expressing Mm523 and revealed with our anti-PGBD5 serum. The top panel shows the nuclear genomic DNA staining by DAPI, the central panel shows the CY3 signal that depicts the distribution of Mm523 within the cell, the bottom panel corresponds to the 22 merged images. These results confirm that the presence of Mm523 within the nucleoplasm is not homogeneous. In all, conclusions for (**C**), (**D**) and (**E**) were identical to those of (**A**) and (**B**).

**Figure S3. Cellular localization by confocal microscopy of PGBD5 into brain cell lines: (A) T98G, (B) H4 and (C) 8-MG-BA.** Top panel: cells revealed with the pre-immune serum. Bottom panels(s): cells were revealed with our anti-PGBD5 serum. The left pictures show the nuclear genomic DNA staining by DAPI, the middle picture show the CY3 fluorescence, the right pictures correspond to merged images. In all, the PGBD5 localization profiles A, B and C revealed its presence both in the cytoplasm and the nucleus of cells. By comparison with the profiles of the PB (10) and *mariner* (37) transposases, the localization of PGBD5 is not exclusively nuclear, its distribution in nuclei being enriched onto the nuclear membrane and not diffuse within the nucleoplasm.

**Supplementary data 1. Sequence of pGH plasmid carrying each one human *pble* and primer sequences for LAM-PCR.**

**Supplementary data 2. Features of chromosomal breakpoint obtained by sequencing LAM-PCR products.**

**Supplementary data 3. RNA-seq profile of our laboratory HeLa cells.**

**Supplementary data 4. Features of ChIP-seq peaks.**

**Supplementary data 5. Annotation of human *pbles*.**

