## Supplementary figures and images for "The *piggyBac* derived transposase 5 (PGBD5) can interact with human *piggyBac*-like elements"

### figures

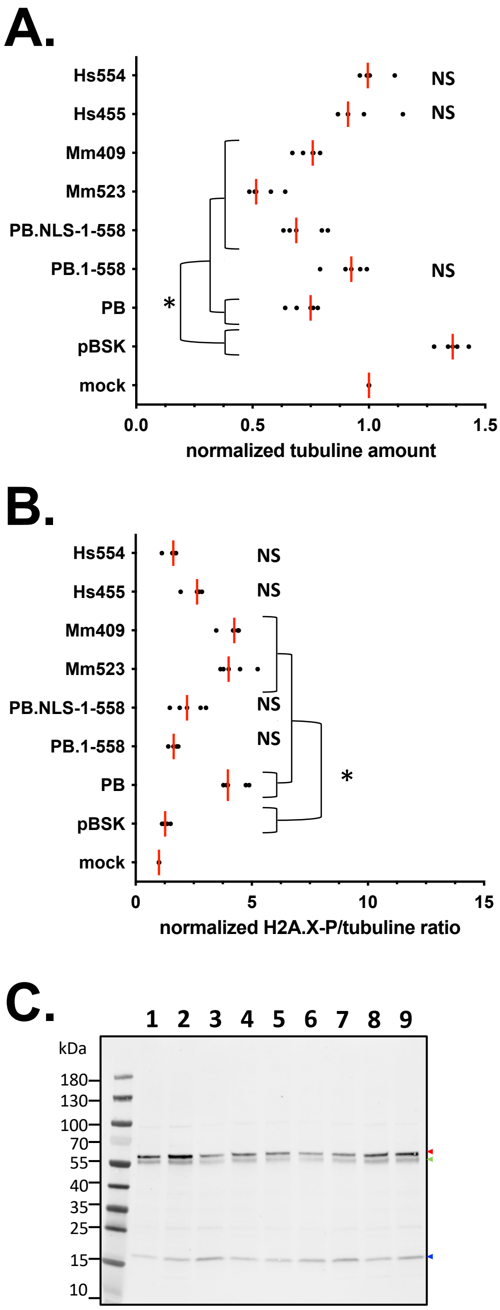

Figure S2A

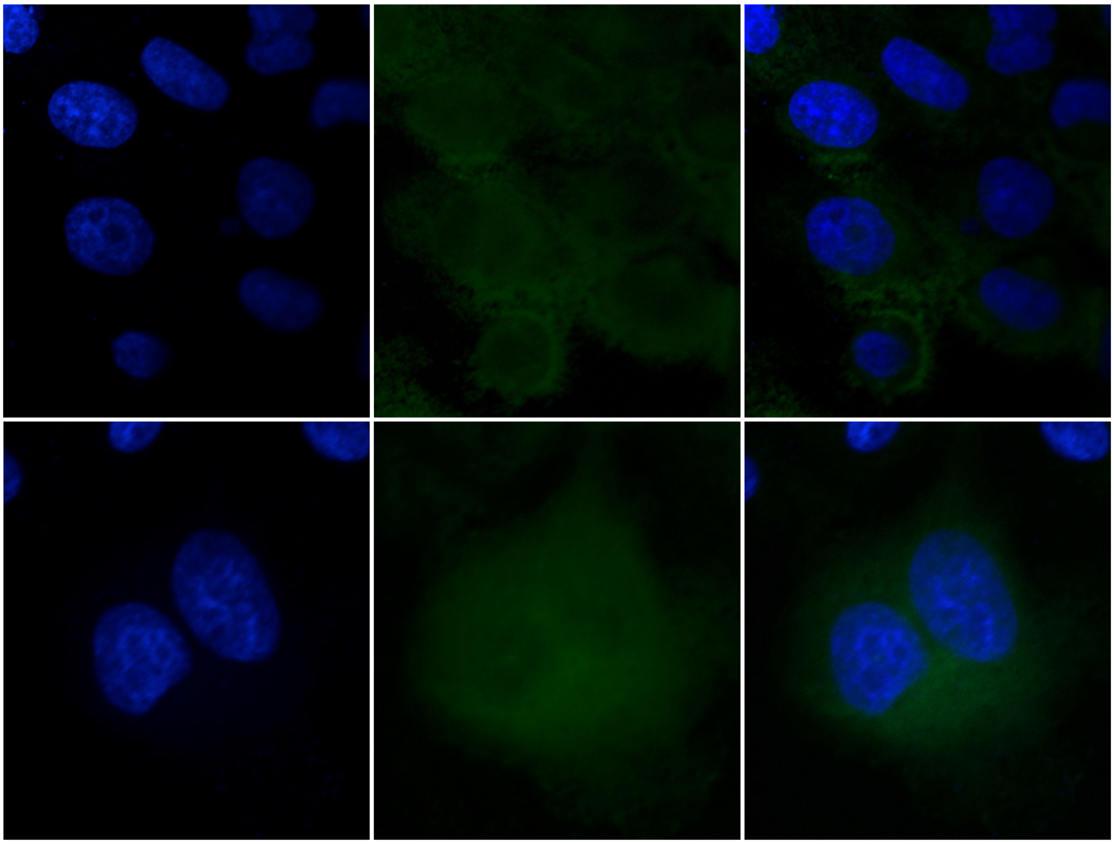

DAPI Mm409-GFP merged

Figure S2B

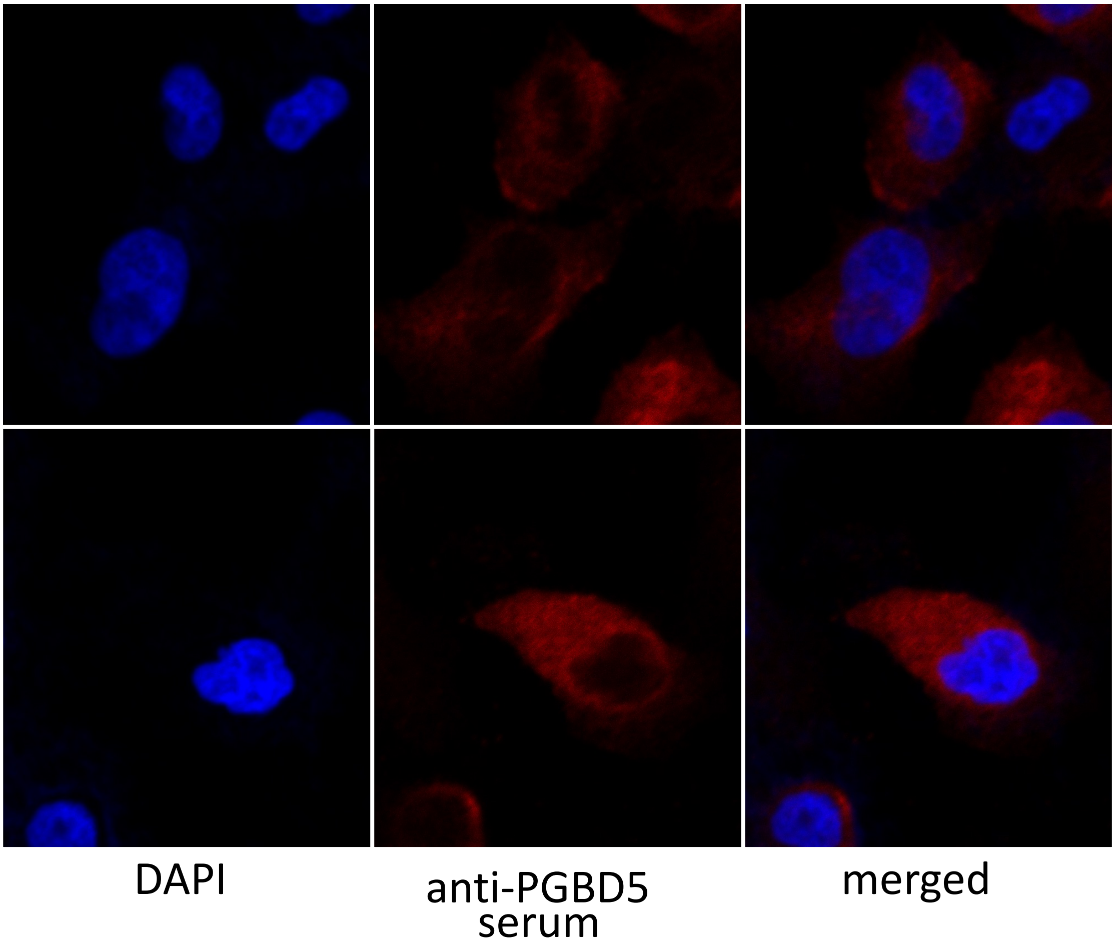

Figure S2C

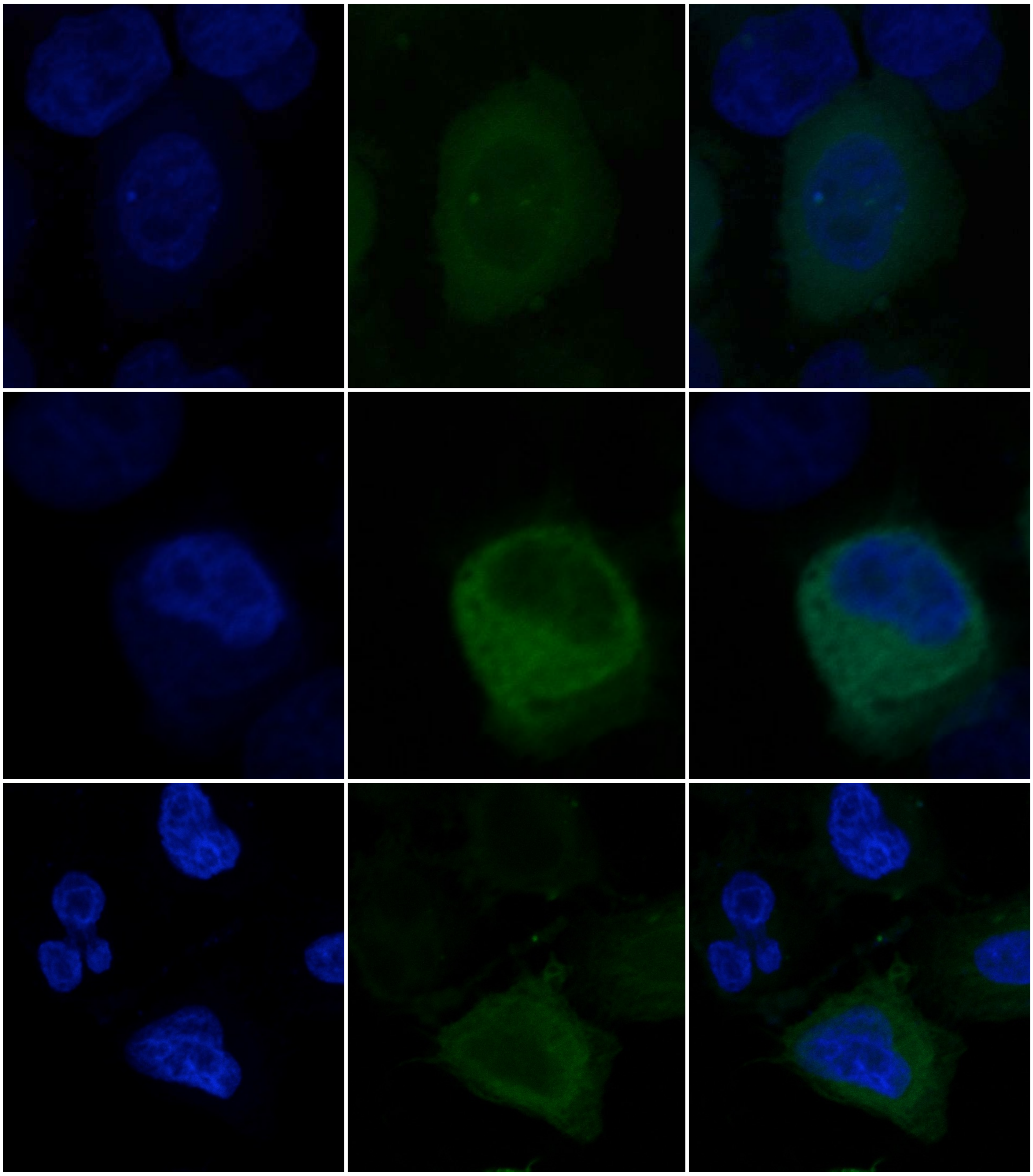

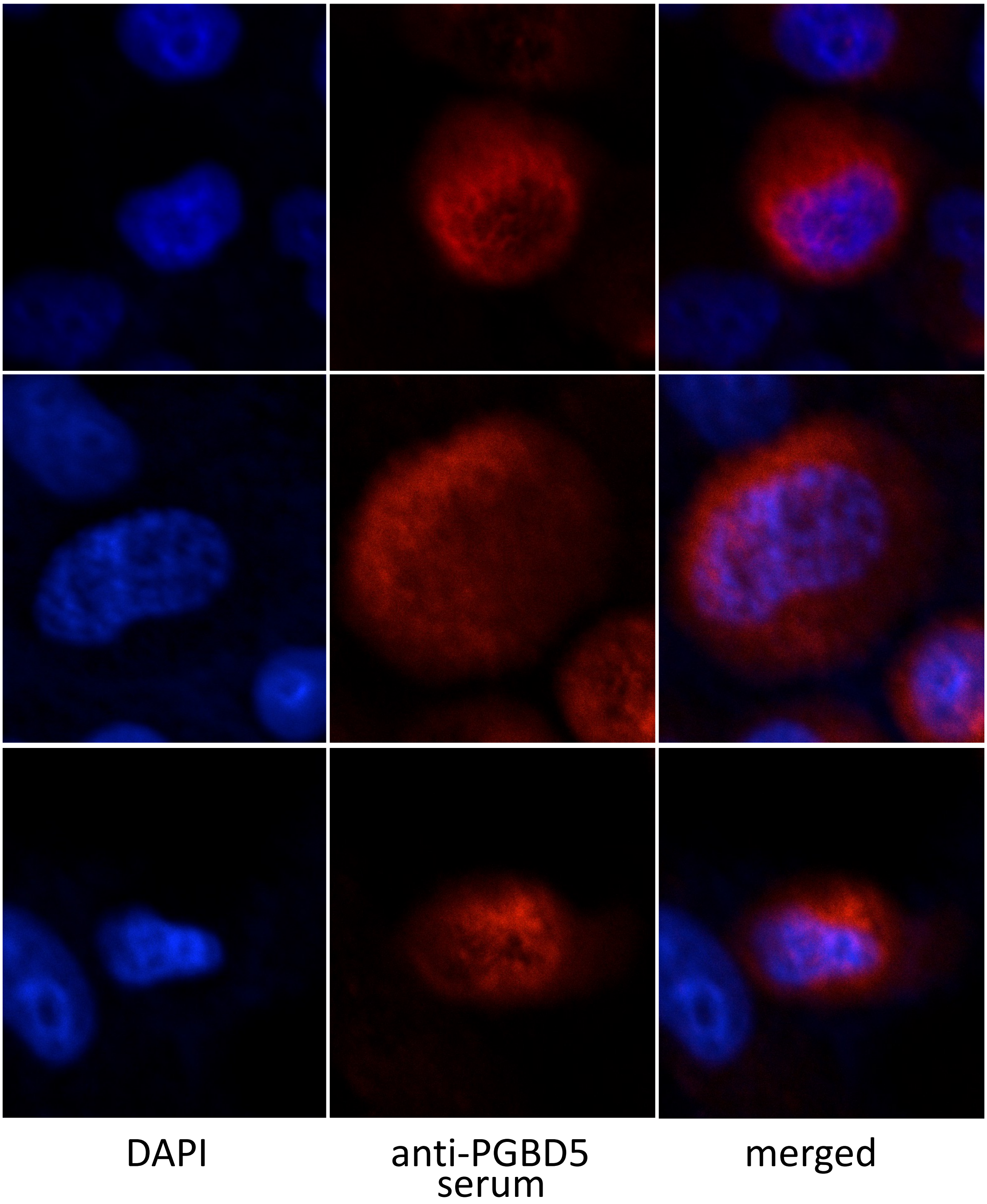

Figure S2E

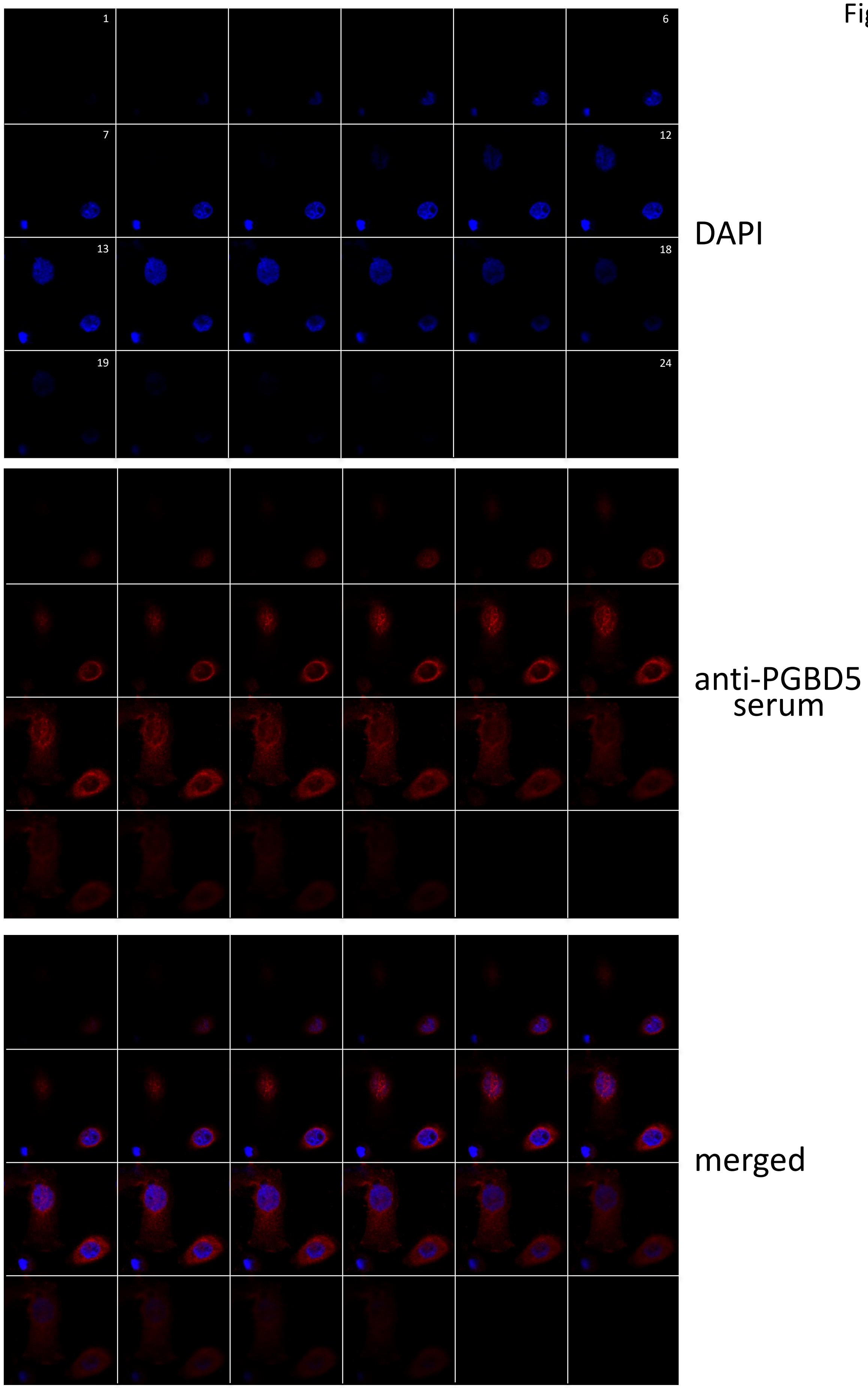

Figure S3A

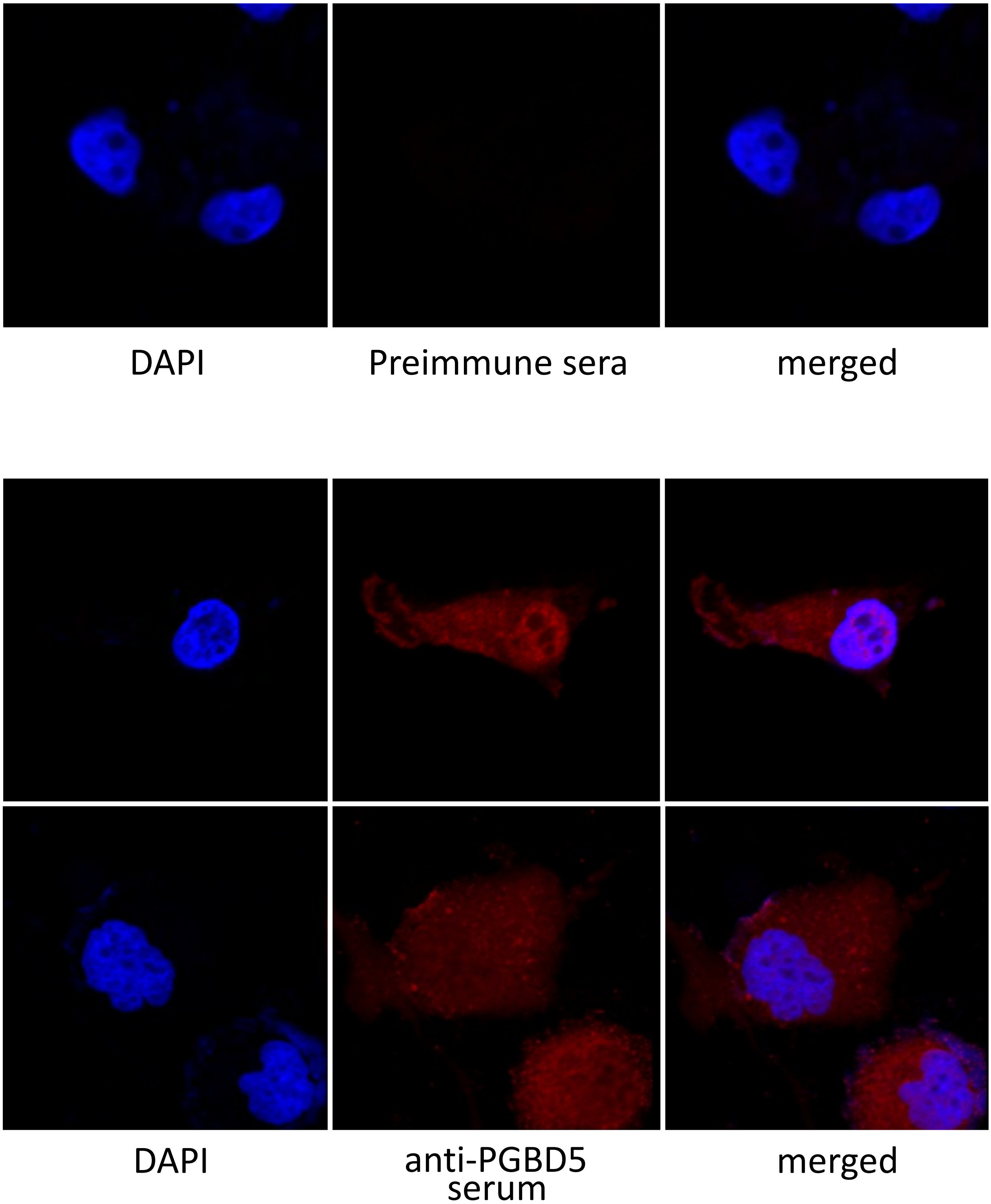

Figure S3B

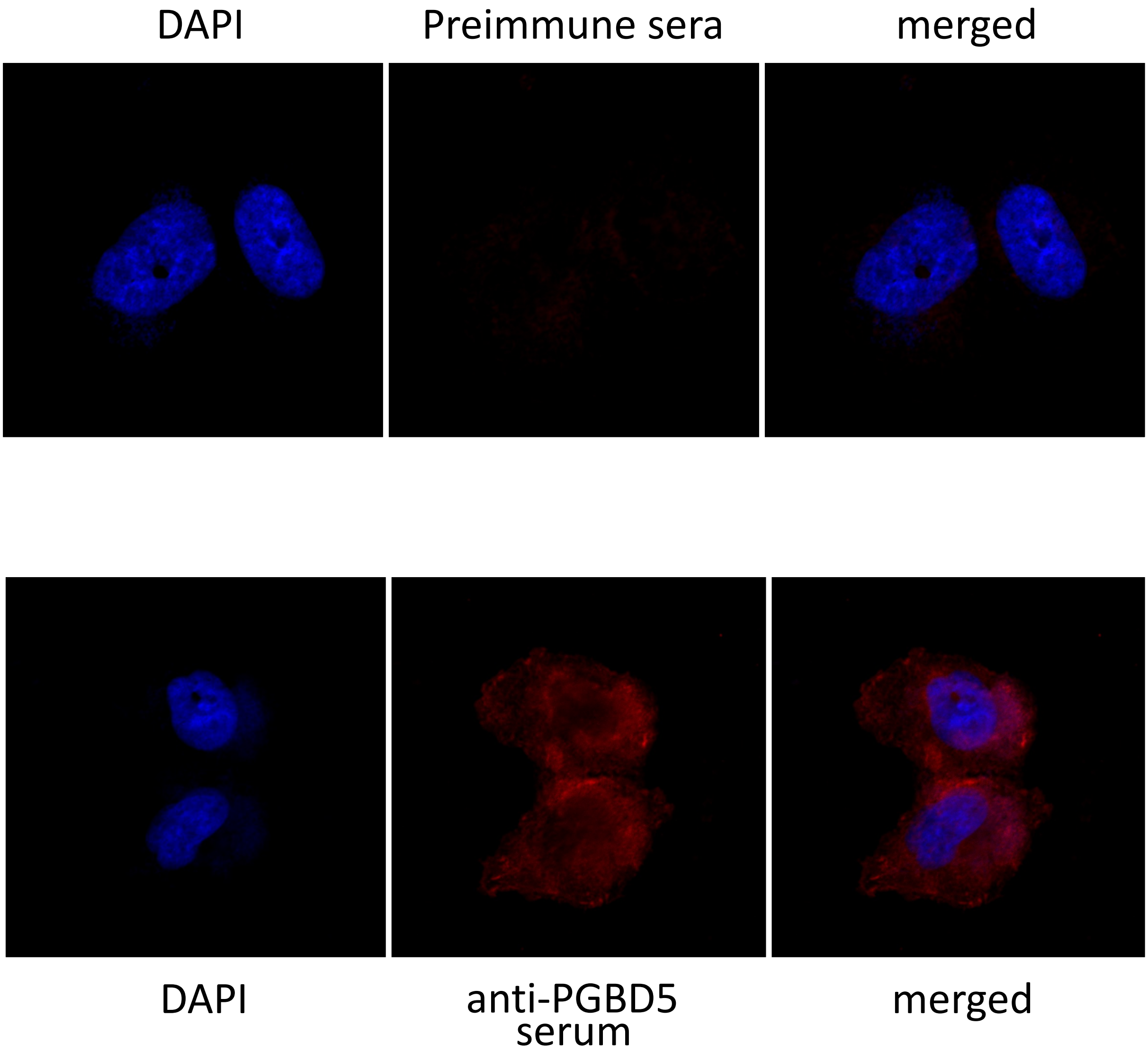

Figure S3C

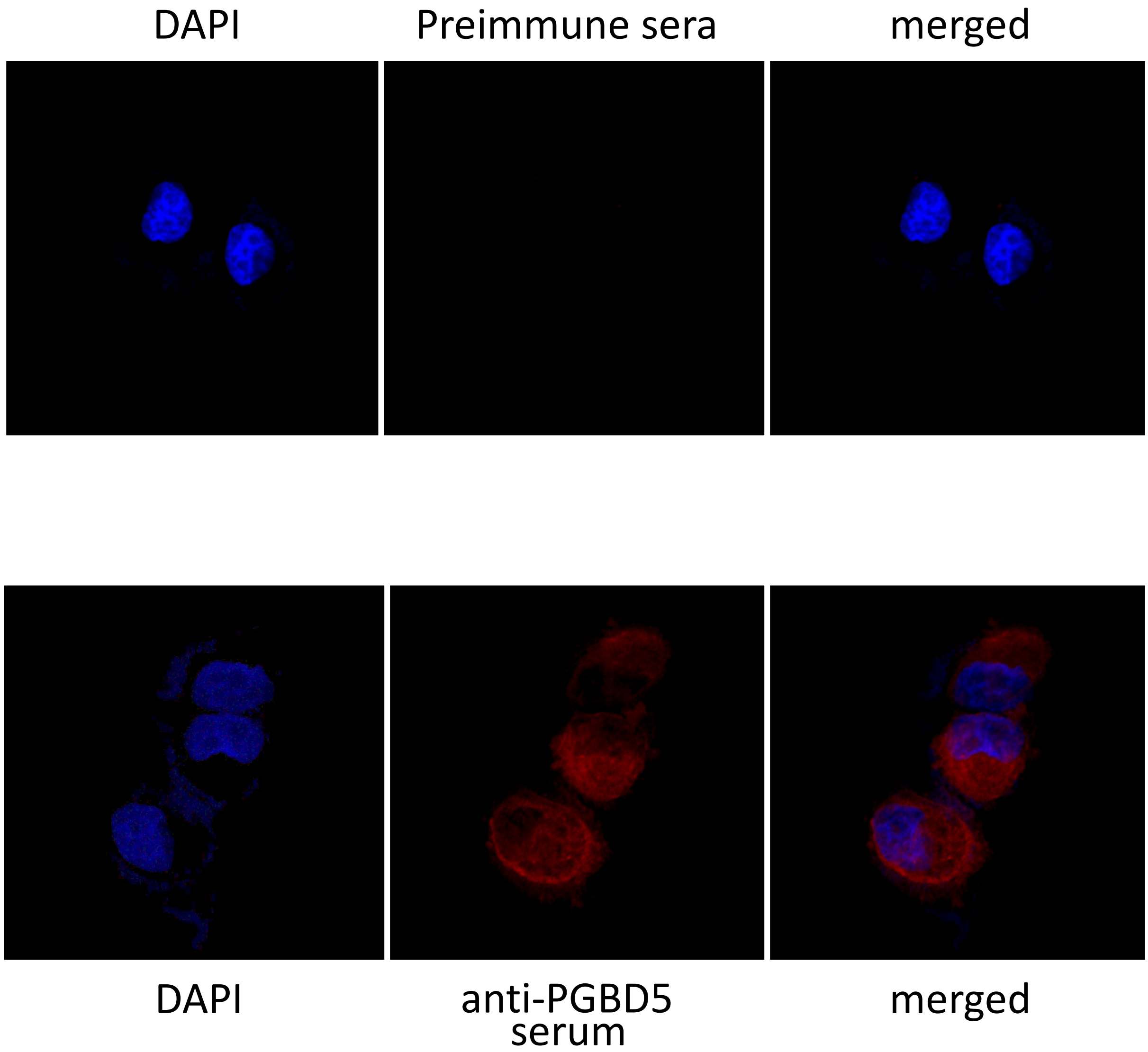
